# The absent P3a. Performance monitoring ERPs differentiate trust in humans and autonomous systems

**DOI:** 10.1101/2025.03.12.642947

**Authors:** Daniel A. Rogers, Kirsty J. Brooks, Anthony Finn, Matthias Schlesewsky, Markus Ullsperger, Ina Bornkessel-Schlesewsky

## Abstract

To address suggestions that human brain responses to autonomous system errors may be used as brain-based measures of trust in automation, the present study asked participants to monitor the performance of either a virtual human or an autonomous system partner performing a novel, complex, real-world image classification task. We predicted visual feedback of partner errors would elicit the feedback-related negativity and P3 ERP components, and that these components would differ between the human and system groups. Behavioural results showed that while participants calibrated their trust in their partner according to our intended manipulation of error rates, no group differences were found. The ERP data, however, revealed FRN and P3 effects for both groups, modulated by accuracy and error rate. An unexpected finding was that the P3 topography differed between groups, with both a frontal P3a and posterior P3b component seen for the human condition, while for the system condition only the posterior P3b was observed and the P3a was completely absent. We suggest that this selective absence of the P3a may reflect reduced frontal attention during system monitoring in passive task conditions potentially resulting from reduced social and emotional processing for the system partner. This study demonstrates the potential for EEG-based measures of trust in automation to surpass the sensitivity of traditional measures of trust, while additionally uncovering a potential neural signature of automation complacency in the absent P3a. This identifies potential boundary conditions under which the application of human correlates of performance monitoring may not apply to the monitoring of an automated system.

---

The rapid uptake of automation and artificial intelligence (AI) into seemingly every domain of human activity, especially complex and safety-critical tasks, has created strong interest in understanding how humans monitor and evaluate these systems. One notable proposal is to use brain responses to observed system errors, specifically feedback-related event-related potentials (ERPs), as real-time, objective measures of human trust in automation (TiA) (Drnec, Marathe, Lukos, & Metcalfe, 2016; Somon, Campagne, Delorme, & Berberian, 2017). This proposal assumes that known neural correlates of human performance monitoring, such as the feedback-related negativity (FRN) and the P3, can be similarly elicited when observing and supervising system performance. This is an assumption which also finds theoretical support in the Computers as Social Actors (CASA) framework, a prevailing view in human-computer interaction and neuroergonomics which proposes that humans treat computers and automated agents as social actors despite knowing they are not human (Nass & Moon, 2000). This view suggests that the social-cognitive processes engaged when monitoring a human may naturally be extended to the monitoring of a system. However, this assumption is currently theoretically and empirically underdeveloped as inconsistent labelling of ERP components is obscuring previous findings while little research has directly compared neural responses to observed human and system performance on the same task.

In the present study we have thus taken a foundational perspective. We aim not only to determine whether the FRN and P3 can be elicited from observed system performance, but also to identify whether any inherent differences in neural responses to observed human and system performance exist by creating a task in which the observed performance is identical across the two conditions. We propose that the observation of a human partner compared to an autonomous system may represent fundamentally different social contexts, and such contextual differences have been shown to modulate ERP responses to observed errors (Koban et al., 2010), with amplitude reductions reported for out-group compared to in-group members (Wang et al., 2017). Further, the human tendency to reduce cognitive engagement when monitoring technological compared to human agents, as seen in well-known phenomena such as out-of-the-loop performance reductions and automation complacency (Parasuraman & Riley, 1997; Endsley & Kiris, 1995), supports the potential for differential processing of human and system performance at a neural level. If such differential processing does exist, this would identify potential boundary conditions under which the direct adaptation of human performance monitoring to system monitoring cannot be directly applied, challenging the assumption of equivalence underlying proposals to adopt neural measures of TiA. In practical terms, this would mean that the use of neural measures of human performance monitoring in system supervision would require establishing the validity of such measures in system-specific contexts prior to their widespread adoption.

To build this foundation, the introduction begins by defining TiA and reviewing current challenges in its measurement, which motivate the need for neural measures of TiA. From a review of performance monitoring components in general, the FRN and P3 are then identified as suitable for the proposed measurement of TiA, highlighting their established role in human error monitoring, identifying their link to expectation violation and trust, and their characteristic elicitation by external feedback. We also clarify that, in our view, several other components elicited by external performance feedback — regardless of the labelling used in prior studies of human trust, human-machine interaction or autonomous system supervision — should be considered feedback-related components, addressing inconsistent labelling in the literature and enabling results from various methodologies to be brought together parsimoniously to support the use of the FRN and P3 as measures of TiA. Finally, we introduce the current study, which uses a complex image categorisation task and systematically manipulates observed partner performance (human vs. system; varying error rates) to test whether FRN and P3 amplitudes vary as a function of trust, partner type, and feedback valence.

## Trust in automation

Automation describes a multitude of technological innovations designed to assist with, and increasingly replace, human task performance (Parasuraman & Riley, 1997). But however capable and efficient technology may be, a human operator is often required to perform a supervisory role and interact with the automation as part of a joint human-automation system (Parasuraman & Riley, 1997). Human-automation systems are intended to improve task performance by utilising the individual strengths of both agents (Drnec et al., 2016). However, humans have historically been poor monitors of automation, which has led to adverse outcomes and death in some cases (Parasuraman & Riley, 1997). Joint human-automation system performance is optimal when the trust a human user has in an autonomous system matches that system’s capabilities, a concept known as Calibrated Trust (Lee & See, 2004). Calibrated Trust is critical to the effective monitoring of automation. If a user’s trust does not match system performance, then states of overtrust or distrust may exist, leading to misuse through over-reliance on autonomy and disuse through inadequate trust in autonomy, respectively (Lee & See, 2004). Measuring extant levels of trust a human user has in an autonomous system is thus crucial to determining Calibrated Trust and the safe, appropriate use of automation.

Measuring trust in automation (TiA) has been shown to be a difficult process, however, as trust is differentially affected by human factors such as attitudes to automation use (Hoff & Bashir, 2015), as well as machine characteristics including competence, predictability and dependability (Merritt & Ilgen, 2008). Importantly these factors can be influential on the real-time performance of humans when making decisions and interacting with automation (Hu, Akash, Jain, & Reid, 2016), meaning that trust levels are not necessarily static and are in fact variable. While a number of previous studies using different tasks such as route planning and visual search have established that automation errors negatively influence user TiA (de Vries, Midden, & Bouwhuis, 2003; Dzindolet, Peterson, Pomranky, Pierce, & Beck, 2003; Lee & Moray, 1994; Moray, Inagaki, & Itoh, 2000), most previous attempts to measure TiA have generally been limited to self-report questionnaires (Drnec et al., 2016). Self-reported levels of trust are not necessarily reliable or objective, and because they are often obtained post-task completion, they do not allow real-time trust in automation to be measured (Drnec et al., 2016). Obtaining an online measure of TiA may enable fluctuations in trust over the course of a task due to human factors and machine characteristics to be identified, allowing steps to be taken to mitigate risks as needed.

### EEG correlates of performance monitoring

Recent reviews have proposed that examining error-related potentials recorded from human observers in response to automated system errors may allow for the development of new brain-based metrics of trust in automation (Drnec et al., 2016; Somon et al., 2017). Error-related potentials follow a reliable pattern of EEG activity associated with performance monitoring and error processing, which can be observed at stimulus presentation, response execution, and during performance feedback (Ullsperger et al., 2014). This pattern is characterised by a fronto-central negativity that is quickly followed by a fronto-central positivity and can be seen for both self-produced and observed errors during task performance and task observation (Ullsperger et al., 2014). This subset of event-related potentials (ERPs) has been utilised for several decades to investigate performance monitoring. Theories of the functional significance of error-related potentials differ, although major accounts suggest that they represent a prediction error signal indicating an unexpected outcome (Ullsperger, 2024).

First identified was the error-related negativity (ERN), also known as the error negativity (Ne), which is a response-locked fronto-central ERP component peaking 80-100ms after errors are committed on speeded reaction time tasks (Falkenstein, Hohnsbein, Hoormann, & Blanke, 1990; Gehring, Goss, Coles, Meyer, & Donchin, 1993). The ERN is considered to be a reliable marker of errors committed during many different tasks that is not stimulus specific (Ullsperger et al., 2014). Co-occurring with the ERN is the Pe, a later positivity (approximately 350ms after response execution) seen over centro-parietal locations (Ullsperger et al., 2014).

Further research has revealed the feedback-related negativity (FRN). The FRN is a modality-independent negativity elicited by external stimuli and peaking from 200-300ms post feedback over fronto-central areas (Ullsperger et al., 2014). Generally seen as a greater negativity for unexpected or negative outcomes, the FRN has been shown to be modulated by factors such as surprise and the magnitude of the error of a predicted outcome (Kirschner, Fischer, & Ullsperger, 2022; Ullsperger et al., 2014). The FRN is generally followed by a P3 (or P300) component with a peak latency between 300-500ms which can have both frontal (P3a) and parietal (P3b) distributions (Ullsperger et al., 2014). Like the ERN, the FRN appears to be robust across different tasks such as time estimation tasks (Gruendler, Ullsperger, & Huster, 2011) and gambling tasks (Ma, Meng, & Shen, 2015; Yu & Zhou, 2006).

More recently, error potential research has been extended to the study of errors committed by others through the discovery of the observed error-related negativity (oERN) and the observed feedback-related negativity (oFRN). Seen on trials where participants observed via a computer screen the errors of a virtual human participant on a choice reactiontime task (participants were told a real human was performing the task in a separate room although they were actually viewing simulated responses based on previous data), the first reported oERN was a negative going deflection peaking at 230ms at electrode Cz (Miltner et al., 2004). The oERN has since been reported in several other studies with a similar topography and latency across different tasks, including the Eriksen flanker task and gambling tasks (Koban, Pourtois, Vocat, & Vuilleumier, 2010; van Schie, Mars, Coles, & Bekkering, 2004; Yu & Zhou, 2006). The oFRN likewise has been reported in studies where participants observed action outcomes of others via computer screen (Bellebaum, Kobza, Thiele, & Daum, 2010; Burnside, Fischer, & Ullsperger, 2019) with a similar topography and latency to the oERN and the FRN. Both the oERN and the oFRN are followed by associated positivities with a latency of 250-500ms post error observation (Koban et al., 2010).

There are similarities between the FRN, oERN and oFRN which are worth noting. All share a similar topography and latency while also being elicited by external feedback rather than being self-generated. Being externally generated may make the FRN and the oERN particularly relevant to the study of TiA, as monitoring an autonomous system is only possible if external feedback is available to the human monitor of said system. Interestingly, the oERN has been suggested to be dependent on a reward or punishment existing for the observer of the task at hand (de Bruijn, Schubotz, & Ullsperger, 2007), whereas the FRN has been reported in tasks where personal consequences of performance did not exist for the observer of the task outcome (Ullsperger et al., 2014).

### The FRN and P3 in system supervision contexts

From the review of components above, the FRN and the P3 emerge as the best candidates for the measurement of TiA in applied scenarios due their robustness across tasks and the fact they can be elicited in passive observation paradigms without personal consequences for the observer of the tasks at hand. This is relevant for real-world human-automation teaming scenarios where humans routinely monitor an automation without directly performing the task themselves. Further, though the adaptation of known correlates of human performance monitoring to system monitoring is not yet well advanced, a range of error-related potentials have been reported during the observation of system errors using a variety of tasks. A review of these studies shows that components sharing the latency and topography of the FRN and P3 have been elicited in supervision contexts previously. While the labelling of components has not followed the FRN and P3 nomenclature, we argue here that these previous findings may be parsimoniously brought together under an FRN and P3 framework which shows that these components can be successfully applied to complex scenarios.

Ferrez and Millán (2008) conducted a human-machine interaction study and reported eliciting what they termed interaction error potentials from participants who were controlling the movement of a simulated robot. Participants tracked robot movement via a computer screen and responses to movement errors elicited an increased negativity peaking 250ms post feedback at electrode FCz compared to correct movements. The negativity was followed by a positivity peaking at 320ms, also over electrode FCz. These components fit the latency and topography of the FRN and P3, and though the authors argued against their findings being feedback-related due to the interaction between the human controller and simulated robot, being generated by visual feedback aligns clearly with known FRN and P3 characteristics. A follow-up study using the same task reported similar components (Chavarriaga & Millán, 2010).

A more recent study utilising a modified arrow flanker task compared observed performance of both humans and systems, and reported finding both an N2 effect and a P3 in response to errors committed by both groups (Somon et al., 2019). The reported N2 effect was maximal at electrode FCz, and the nature of the eliciting stimulus being visually presented feedback of the partner responses suggests this could be considered an FRN followed by a corresponding P3. Notably, this study reported that the P3 amplitude was significantly reduced for the observed system errors compared to human errors, a result which demonstrates the need to compare system and human performance on the same task when considering the adaptation of correlates of human performance monitoring to TiA. It is relevant to highlight here that participants in this study were only allowed 10ms to perceive the target themselves, so their judgements of partner accuracy for each trial were not necessarily certain. This is important to note as the FRN and P3 classically are generated when outcomes are known, and any uncertainty as to the correct response may obscure or attenuate them in this type of paradigm.

### The FRN and P3 in trust contexts

There is a widely held view that the FRN is a neural index of expectation violation that reflects deviation of outcomes from prior predictions (Kirsch et al., 2022; Ullsperger et al., 2014). This is directly relevant to the measurement of trust, as trust is in effect a calibrated expectation about the future performance or behaviour of another person or agent (Lee & See, 2004). On this basis, the FRN may provide a real-time measure of trust violation as when a highly trusted partner produces an unexpected error, this violation of expectations should be observable in FRN amplitudes.

Consistent with this reasoning, previous work has found that the FRN may be used as an online index of human-to-human trust. Ma and colleagues (2015) found that trust outcomes for decisions to trust a human partner were reflected in FRN amplitudes in a modified gambling task, with increased amplitudes seen for rewarded trust decisions. However, the P3 results of this study were unfortunately not reported. An earlier study by Long, Jiang, and Zhou (2012) similarly reported that, in a modified coin-toss game, correct decisions to trust others elicited increased FRN amplitudes compared to correct decisions not to trust. The FRN for rewarded trust decisions in this case was however not found along with an increased P3 component, differing from the classical pattern of performance monitoring activity. In both studies, participants were convinced that they were interacting with human partners when performing the tasks. However, this was not the case and all facets of the tasks completed were presented via a computer screen in a predetermined and scripted manner. It is interesting to consider how these results may apply to the study of TiA, for while the study methods were highly reliant on computer scripting, suggesting a similarity to the operation of an automated system, how the participant responses may have differed had they known they were interacting with a computer script rather than a supposed human is not known. Further to the results of Somon et al. (2019) reported above, this also highlights the need to consider psycho-social differences between observing human and system groups and how this may influence the observed ERP results.

One recent study by de Visser et al. (2018) has directly investigated the possibility of using error-related potentials as an index of human TiA. The authors reported an oERN and oPe in response to automated system errors on a modified flanker task. Most interestingly, the authors were able to show that subjective ratings of trust in automated system task performance were a significant predictor of the oPe, with increased trust related to increased oPe amplitudes. However, this relationship was limited to the oPe as a similar effect of trust ratings on oERN amplitudes was not found. Following previous work on the oERN in human observation (van Schie et al., 2004), the target stimulus was presented to observers without the distractor arrows present to minimise the uncertainty of the required correct response. The eliciting stimulus in this experiment was again visually presented feedback indicating the automated system’s chosen response. The authors of the study also manipulated the accuracy of the automated system (a measure of system reliability) by using two levels of performance – high (90% correct) and low (60% correct), respectively – that alternated between blocks and which were related to subjective measures of trust with increased trust related to high accuracy. Increased oPe amplitudes were found for less frequent errors in the high performance rate blocks as compared to the more frequent errors in the low performance rate blocks, however it was also found that performance was quite predictable, and participants quickly recognised the performance level of the system for a given block. The authors suggest this may account for the failure to relate the oERN in this study to the subjective measures of trust. The oERN and oPe reported in this study could also plausibly have been characterised as an FRN and P3, as they were induced by visually presented feedback.

The studies discussed above have shown that the FRN and P3, and components sharing their characteristics, can be observed in response to both simple and more complex visual feedback during the observation of automated systems. In addition, these components have been reliably elicited by relatively simple paradigms such as the Flanker task as well as more complex tasks including simulated human-machine interaction and gambling tasks. The finding of an FRN and P3 using different visual feedback characteristics and different tasks suggests the possibility that these components may be well suited to applications in more complex real-world environments where automated systems are in common use. However, despite this robustness of the FRN and P3 (and similar components) across tasks, studies have shown mixed results for observed partner type, with an increased FRN found when trusting a human by Long et al. (2012) and an oPe seen for increased trust in the study of TiA by de Visser et al. (2018), while Somon et al. (2019) reported increased P3 amplitudes to observed human compared to system errors. These discrepant findings for partner type indicate further research is required to establish how brain responses to observed partner types may differ on the same task before neural measures of TiA can be utilised in real-world environments.

### The present study

To investigate the potential use of the FRN and P3 as brain-based indexes of human trust in autonomous systems, we created an experimental task that extends beyond the traditional paradigms and feedback used in FRN research into a more real-world environment. Our paradigm utilised a complex image recognition task inspired by the Asirra CAPTCHA (Elson, Douceur, & Howell, 2007), an image categorisation task which is simple for humans to perform but which is presumed to be difficult for automated algorithms to master. Participants acted as monitors of either a (virtual) human or an automated system partner who performed the task. In reality, both partners were pre-scripted and observed performance was identical across partner types. The FRN and P3 were time-locked to the onset of visual feedback given to participants on each trial to indicate the partner’s chosen image. The observed performance of the human or system partner fluctuated between one of 12 separate error rates, in an ordered but unpredictable manner, thus allowing us to examine the effects of differing performance levels while avoiding the confound of participants guessing the upcoming performance. Participant trust ratings in their partner were given at regular intervals and could thereby be compared to FRN and P3 amplitudes to observed performance feedback.

Specific aims of the study were to identify how observed partner performance, partner type (human and autonomous system), and feedback valence (error and correct) affect reported participant trust ratings (performance predictions) and recorded neural responses to partner performance feedback in a complex real-world task. We tested the following hypotheses. **H1**: Observed partner performance will influence participant trust ratings (performance predictions), with high error rate blocks associated with worse predictions of performance than medium or low error rate blocks, as supported by de Visser et al. (2018), who demonstrated that automated system error rates directly influence subjective trust ratings. **H2**: Visual feedback of partner error trials in a real-world complex image categorisation task will induce increased FRN and P3 amplitudes compared to correct trials. This is supported by the proven robustness of the FRN and P3 across multiple paradigms. **H3**: P3 amplitudes will be increased for human partner errors in comparison to system partner errors as found by Somon et al. (2019), who directly compared responses to human and system on the same task. We further explored the effects of partner performance (error rate), trust in partner (performance predictions), and partner type (human or autonomous system) on observed FRN and P3 amplitudes to extend previous findings to a complex real-world task with unpredictable error-rate variance, which is a previously unexplored experimental context.

## Methods

### Participants

Forty-eight adult volunteers (age range: 18-39 years; mean 25.5; 34 female) participated in this experiment after providing written informed consent. Participants were all righthanded with normal or corrected to normal vision and no history of reading, neurological or psychiatric disorders. Twenty-four participants (age range: 18-39 years; mean 26; 18 female) were randomly allocated to monitor a virtual human partner, with the remaining twenty-four participants (age range: 18-39 years; mean 25; 16 female) randomly allocated as monitors of an autonomous system partner. Participants received a $40 honorarium at the end of the experiment. The research was approved by the University of South Australia ethics committee. Sample size was determined using comparable EEG studies investigating feedback-related ERP components. Burnside et al. (2019), Somon et al. (2019), de Visser et al. (2018), and Ma et al. (2015) all detected FRN and P3 effects (or similar components) with samples ranging from 17-21 participants per condition. Our sample of 24 participants per condition exceeds these precedents, while our use of single-trial analysis methods enhances sensitivity to effects, as practiced by Burnside et al. (2019).

### Design

This study used a repeated measures between-groups design to investigate the effect of partner type (virtual human, autonomous system) and partner error rate (low, medium, high) on error-related brain activity and behavioural monitoring responses to partner performance on a novel image classification task. Participants were randomly allocated to one partner condition, and all error conditions were counterbalanced across participants. In the virtual human monitoring condition, participants were advised that a colleague of the researchers was seated in a nearby room and performing the image classification task remotely. Participants monitoring the autonomous system were advised that they would monitor the performance of a deep learning algorithm that had been trained to classify images.

## Materials

### Task

The task used in this experiment was inspired by the Asirra (Animal Species Image Recognition for Restricting Access) CAPTCHA (Elson et al., 2007). Asirra utilised images of cats and dogs as stimuli for categorisation by either a human user or an automated classification system. Asirra was presumed to be secure as the classification of cats and dogs is a task that is easy for humans to perform which has also been shown to be difficult for an automated classifier to achieve similar performance (Elson et al., 2007).

### Task design

In our task, participants monitored their respective partner type perform the image classification by watching a computer screen and observing their partner’s image selections. The task required the partner to find and select a target image of a dog. On each trial, four images were presented simultaneously in one of four locations (top left, top right, bottom left, bottom right), always with one dog present along with three distractor images of cats. The location of the target image was randomised across trials to replicate the way a real CAPTCHA varies target locations. On presentation of the four images, participants were given 1600ms to find and fixate on the target image themselves, after which the images remained on-screen while a yellow border highlighting the partner’s selected image was given as feedback for 800ms. Participants were not required to give any kind of response. The full trial sequence was as follows: blank screen (300ms), fixation cross (600ms), target and distractors displayed (1600ms), partner feedback (800ms), (see Figure 1). Participants completed two training blocks of fifty trials in which the partner error rate was set at 20%. The actual experiment consisted of 600 trials across twelve experimental blocks of between forty-seven to fifty-three trials with between five to sixteen errors in each block. For both partner conditions, participants were advised that their respective partner was required to make the correct selection on approximately 80% of the trials they observed in order to meet performance expectations. Additionally we advised participants that they could expect their partner to perform at this level. To account for the human partner committing errors on what was apparently an easy task for humans to perform, participants were advised that the human partner was put under pressure by having only half of the time they themselves had to find the target (800ms), while also needing to press a corresponding keyboard button for their selected image based on the location on the screen. Monitors of the autonomous system were advised that the image classifier was trained on a set of 100 sample images and then applied this training to the test images during the experiment.

**Figure 1.**
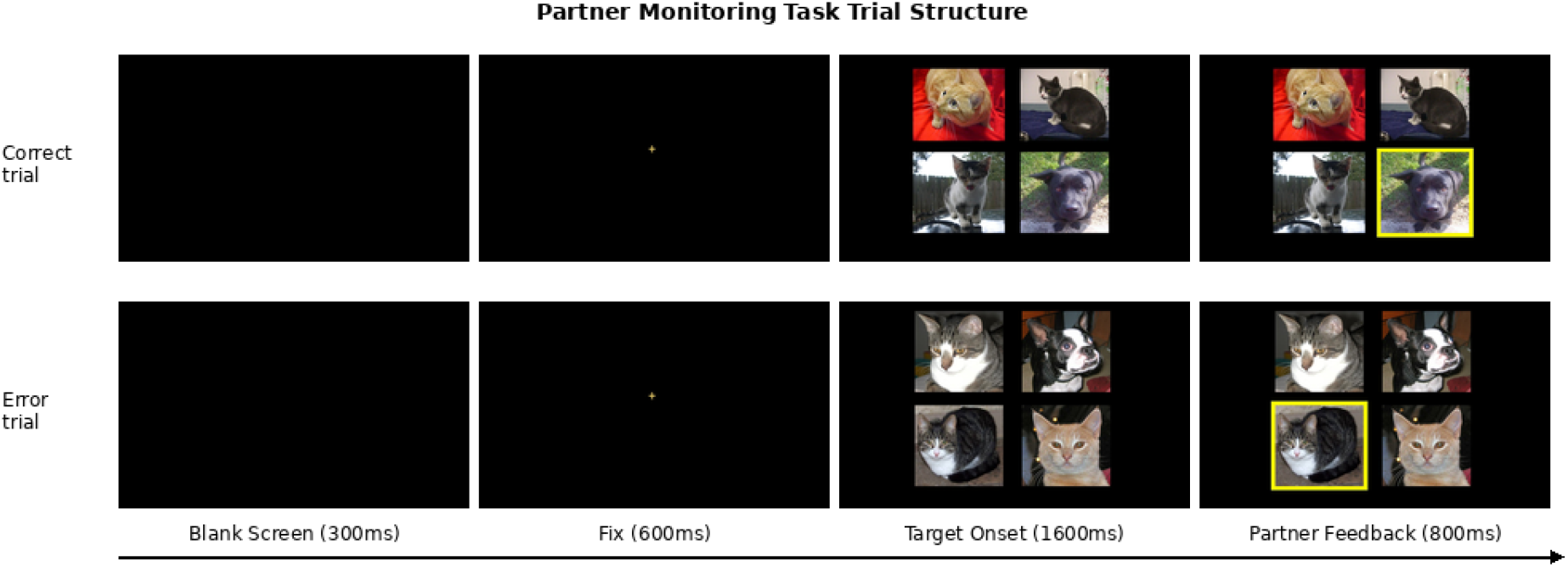
Partner monitoring task trial structure. Trials began with a blank screen and fixation cross, after which a target array was presented. Participants searched for and located the target dog themselves during the 1600ms target onset stimulus period, after which the partner feedback indicated the partner made either a correct (target dog selected; top) or incorrect (distractor cat selected; bottom) identification.

### Image and feedback stimuli

The dog and cat image stimuli were sourced from the Microsoft Kaggle Dogs vs. Cats dataset (Microsoft, 2017). The Kaggle dataset consists of 25,000 unique images of dogs and cats that Microsoft has made available for research purposes. From these images we selected a subset of 2100 cats and 700 dogs. Images were selected according to the following criteria: only one animal per photo; no humans were visible; animal’s head and face were visible; image was in landscape orientation with between a 70-80% height to width ratio. Images were resized with height and width dimensions of 300 pixels by 400 pixels. Image luminance was determined using the PIL (Umesh, 2012) package in python and luminance values were used to sort images to reduce brightness variations in each block.

### Partner error rates

In order to reduce the predictability of partner performance, a series of randomisations of the error rates and block lengths was performed. First, the error rates of five to sixteen errors per block were randomly matched twice to a block length. This resulted in two lists of twelve randomly matched error rates and block lengths. The error rates were then binned in thirds as either low (5,6,7,8), medium (9,10,11,12) or high (13,14,15,16) error rates and for each of the two lists, two randomisations of each third were created (e.g. 6,8,5,7 and 5,8,7,6). For each list the randomised thirds were then used to create block orders which progressed in thirds from e.g low/medium/high. All possible combinations of thirds were created for each list twice (see Supplementary Figure A1), resulting in twelve randomisations of binned error rates per list.

### Performance and trust ratings

Participants were asked to rate their partner’s performance at the end of each block by indicating if they were meeting, exceeding, or falling short of performance requirements. If the partner had achieved the approximately 80% correct required performance rate then participants were to rate them as having performed “as required”. If participants judged their partner had been correct on more than 80% of trials then they were to rate them as having performed “better than required”. Finally, a judged performance of less than 80% correct trials was to be rated as performing “worse than required”. Participants were advised to use the training blocks which were set at 80% correct as a guide for judging the experimental blocks. Participants were deliberately not advised to count errors during the task.

To measure participant’s initial expectations for their partner’s performance, we asked them to predict how their partner would perform in the first experimental block using the same measure as the actual performance ratings. To measure participant’s ongoing expectations and trust in their partner’s performance, at the end of each experimental block they were asked to make a prediction for the following block based on the performance seen so far, again with the same rating options. These participant responses were assigned scaled values (as required = 0, better than required = 1, worse than required = -1) to allow an average rating score across the experiment and error rates to be calculated. To track the participant’s ability to accurately monitor their own predictions, we asked them after each block if the partner’s performance had matched their previous performance prediction with a yes or no response required.

### Procedure

After giving informed consent participants spent several minutes responding to personality, sleep, dispositional trust, and attitudes to automation questionnaires. For EEG testing participants were alone, comfortably seated in front of a computer screen, with a keyboard and mouse used to give all required behavioural responses. After being advised of the format of the experiment, participants read initial instructions for their task on screen and completed the two practice blocks which took approximately 8 minutes. The researchers confirmed after each practice block that the participant was able to locate and fixate on the target images on each trial and that they were able to track the 80% performance example. After confirming participants understood the task, participants were left alone to complete the main experiment which began with a repeat of the instruction on screen. It took approximately 48 minutes to complete all 12 experimental blocks. Following the experimental task participants completed an Eriksen flanker task (not reported here). Resting state EEG recordings were taken for two minutes at the beginning and end of the testing session.

### EEG Recording and Preprocessing

EEG was recorded using a BrainAmp DC amplifier at a 500Hz sampling rate from 32 sintered Ag/AgCl electrodes mounted in an elastic cap (Easycap GmbH, Herrsching, Germany) per the international 10-20 system. Bi-polar electrooculograms (EOG) were recorded via a BrainAmp ExG amplifier, with the horizontal EOG recorded from the outer canthus of each eye and the vertical EOG recorded from above and below the left eye. Ground and reference electrodes were located at AFz and FCz. Electrode impedance values during recording were aimed to be below 20 kOhm.

Pre-processing of EEG data was completed using MNE Python version 0.20.5 (Gramfort et al., 2013) with further utility functions taken from the philistine package (Alday, Appelhoff, & Bornkessel-Schlesewsky, 2023). Correction of EOG artefacts was performed using Independent Component Analysis (ICA). To achieve this, a bandpass filter from 1 to 40 Hz was applied to the raw data and then individual independent components were identified using the FastICA method for the EEG channels only. Epochs were rejected if the peak to peak voltage exceeded 250 microvolts. EOG events were identified using the MNE function “create_eog_epochs”. The function “ica.find_bads_eog” was then used to remove any independent components that were correlated to the EOG activity from the raw data. Following the ICA, data were bandpass filtered from 0.1 to 30 Hz to remove any slow drifts or higher frequency noise. Data were re-referenced offline to a common average reference, and the online reference channel FCz was re-instated by interpolating surrounding channels. Epochs with a 200ms baseline correction were created from -200 to 800ms relative to the presentation of partner image choice feedback. Epochs were rejected if peak-to-peak amplitudes for EEG channels exceeded 150 microvolts while epochs were identified as flatlining and removed if peak-to-peak amplitudes were below 5 microvolts.

### Data analysis

All statistical analyses were performed in *RStudio* (Posit team, 2023) using *R* Version 1.4.1106 (R Core Team, 2023). For data importing and manipulation the *tidyverse* package v.2.2.0 (Wickham, 2023) was used. Modelling of the data was performed using generalised linear mixed effects and linear mixed effects models produced with the *lme4* package v.1.135.1 (Bates, Mächler, Bolker, & Walker, 2015) while *p* value estimates for all effects were obtained from type II Wald χ^2^ tests using the *car package* v.3.1-2 (Fox & Weisberg, 2019). Data plots were created using the *cowplot* v.1.1.1 (Wilke, 2020), *effects* v.4.2-2 (Fox & Weisberg, 2019) and *ggplot2* v.3.4.4 (Wickham, 2016) packages. Model output tables were produced with the *lmerOut* v.0.5.1 (Alday, 2021) and *kableExtra* v.1.3.4 (Zhu, 2021) packages. Contrasts for categorical predictor variables in the statistical models used treatment coding. Baseline levels for the contrasts were assigned as follows: accuracy condition (correct), group condition (human), error rate condition (medium or “as expected” rate), pre-block prediction (medium or “as expected” rate). Error bars in all plotted model figures show 83% confidence intervals which are significant at the .05 level when non-overlapping.

### Behavioural Data

Behavioural analysis used a generalised linear mixed effects model to investigate the fixed effects of error rate and group on the dependent variable monitoring accuracy with random intercepts by participant ID. A linear mixed effects model was used to investigate the same fixed effects also with random intercepts by participant ID for the pre-block performance prediction data.

### EEG Data

The following electrodes were identified as a subset of interest and used for all EEG analyses: F3, F4, C3, C4, P3, P4, FCz, Fz, Cz, Pz, FC1, FC2, CP1, and CP2. We identified the maximal FRN, P3a and P3b effects for the EEG analyses based on the method used in Burnside et al. (2019) where the FRN was defined as the most negative going peak at FCz between 200-350ms, and both the P3a and P3b identified as the most positive deflections between 350-500ms at FCz and Pz, respectively. Based on visual inspection of the grand-average waveforms, we selected time windows for analysis of 190-280ms for the FRN and 350-500ms for the P3. All EEG analyses were conducted on single trial data using linear mixed effect models. Fixed effects included in the models were accuracy, group, error rate, pre-block performance prediction, and sagittality. The sagittality effect was included as a continuous predictor to allow more in-depth analysis of the topographical differences in the data and was implemented using positional co-ordinates available from http://robertoostenveld.nl/electrodes/besa_81.txt. The models also included random slopes and intercepts for accuracy and participant ID.

## Results

### Behavioural Data

#### Partner Monitoring Accuracy

Judgements of partner performance made by participants at the end of each block of trials were identified as correct if participants accurately classified the observed performance rate. These data were used to obtain a measure of partner monitoring accuracy which is visualised in Figure 2A as a raincloud plot by ID, group & error rate. The plotted data represents the overall proportion of correct responses during the monitoring task, with a score of 1.00 representing 100% accurate judgements by a participant for all four observed blocks of a specified error rate. It can be seen that partner monitoring accuracy was consistent across groups, while monitoring of the medium error rates appears to have been the most difficult as only 1 participant managed to correctly rate performance in all medium error rate blocks. Statistical analysis of the partner monitoring accuracy data performed using a generalized linear mixed effect model revealed a significant effect of error rate [type II Wald test: *χ*^2^(2) = 17.20, *p* < 0.001]. No effect of group or interaction of group by error rate was found (see Supplementary Table A1). The modelled data plotted in Figure 2B (refer to Supplementary Table A2 for the full model summary) shows the effect of error rate with medium error rate blocks being monitored less accurately than either low or high error rate blocks.

**Figure 2.**
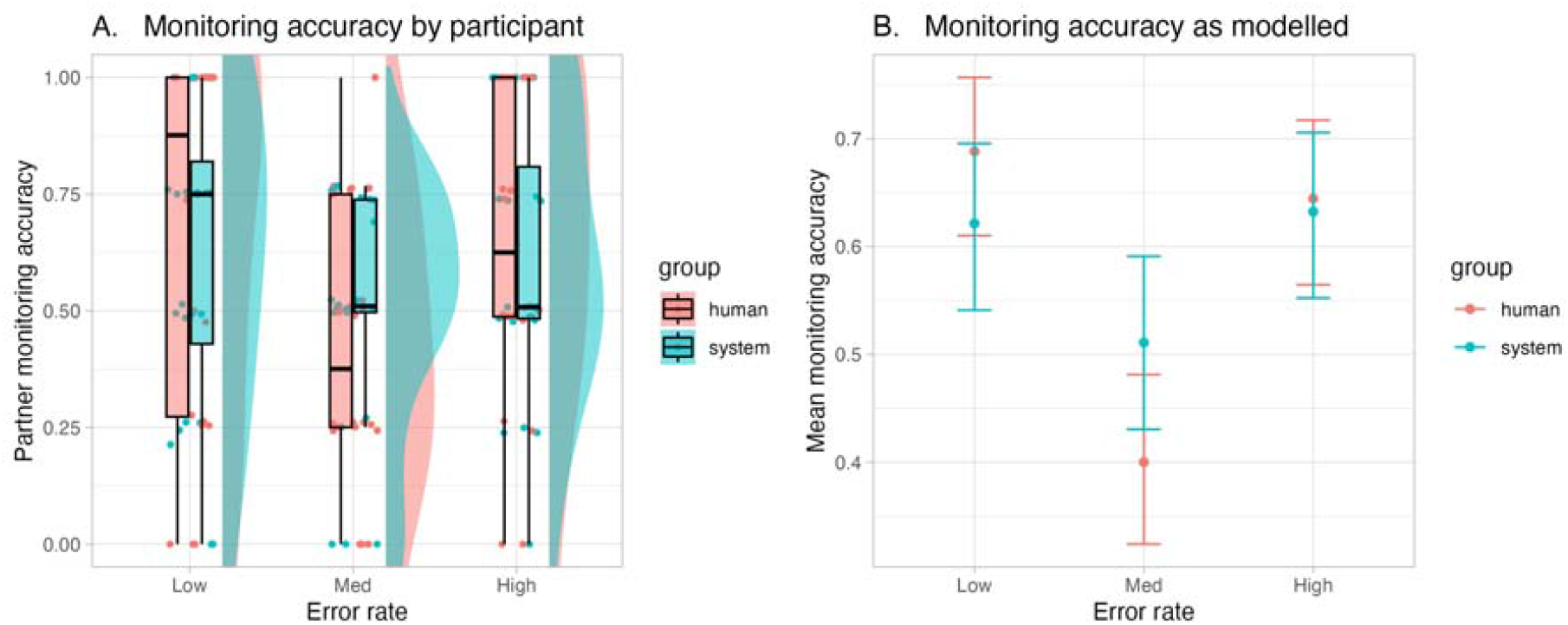
(A) Raincloud plot of Monitoring Accuracy for the partner observation task by Group and Error Rate. Individual participant data points represent by participant accuracy for each error rate condition. (B) Modelled Monitoring Accuracy data for the partner observation task by Group and Error Rate. Error bars represent 83 percent confidence intervals.

#### Performance Predictions

The scaled values for participant’s pre-block performance predictions are visualised in Figure 3A as a raincloud plot by ID, group & error rate. There is a clear trend across both groups for average predicted performance to be worse as error rates increase. The statistical analysis of the predicted performance data again showed a significant effect of error rate [type II Wald test: *χ*^2^(2) = 41.90, *p* < 0.001] with no effect of group or interaction between group and error rate found (see Supplementary Table A3). The modelled data visualised in Figure 3B (see Supplementary Table A4 for the full model summary) shows that predicted performance was best for the low error rate blocks with predictions worsening as error rates increased.

**Figure 3.**
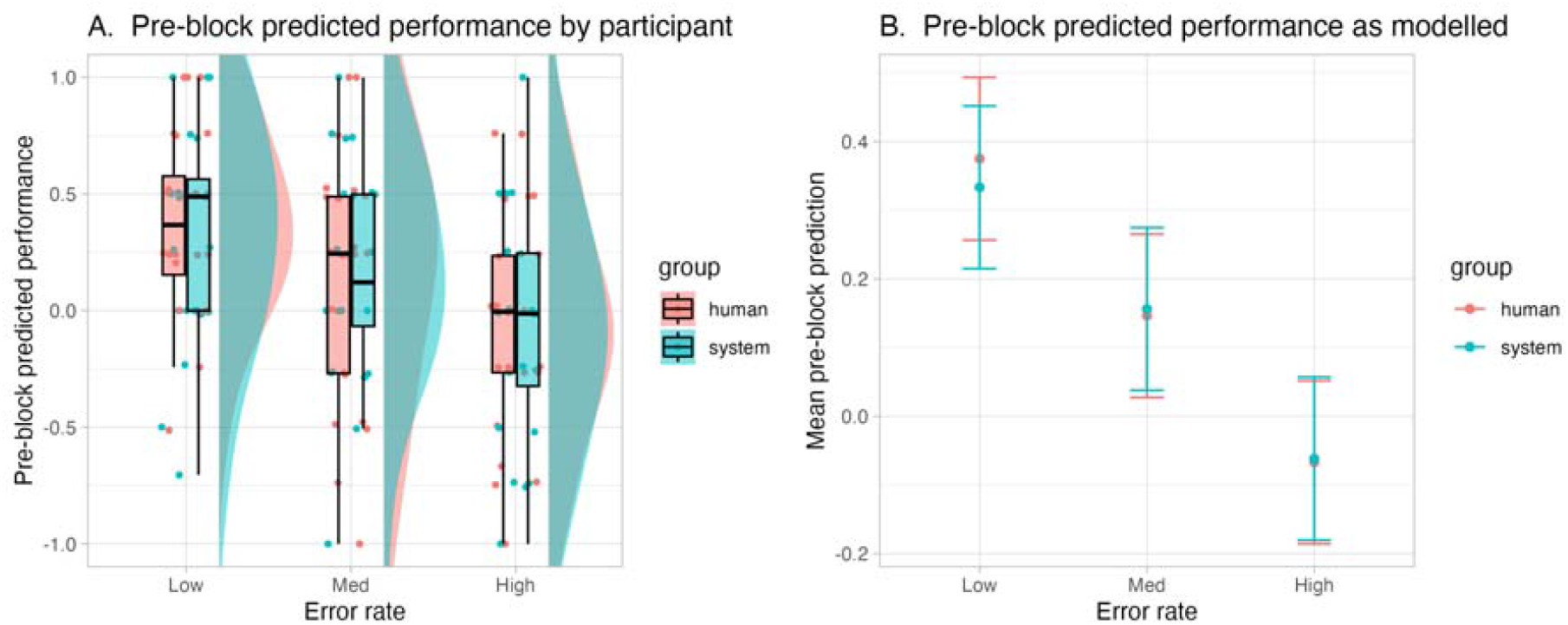
(A) Raincloud plot of Pre-Block Performance Predictions by Group and Error Rate. Individual data points by participant represent mean pre-block predictions for each error rate condition. (B) Modelled Pre-Block Performance Predictions by Group and Error Rate. Error bars represent 83 percent confidence intervals.

#### ERP Data

Grand average ERPs time-locked to critical event onsets for errors and correct partner feedback trials are shown in Figure 4. An FRN effect can be seen for both the human and system groups at electrodes FCz and Cz, with errors producing more negative amplitudes than correct trials. For the human group a P3 effect can be seen over electrodes FCz, Cz and Pz with increased amplitudes seen for error trials. There is also a P3 effect for the system group; however, the topography differs from the human group, with the effect not being observable over electrode FCz and only appearing over electrodes Cz and Pz.

**Figure 4.**
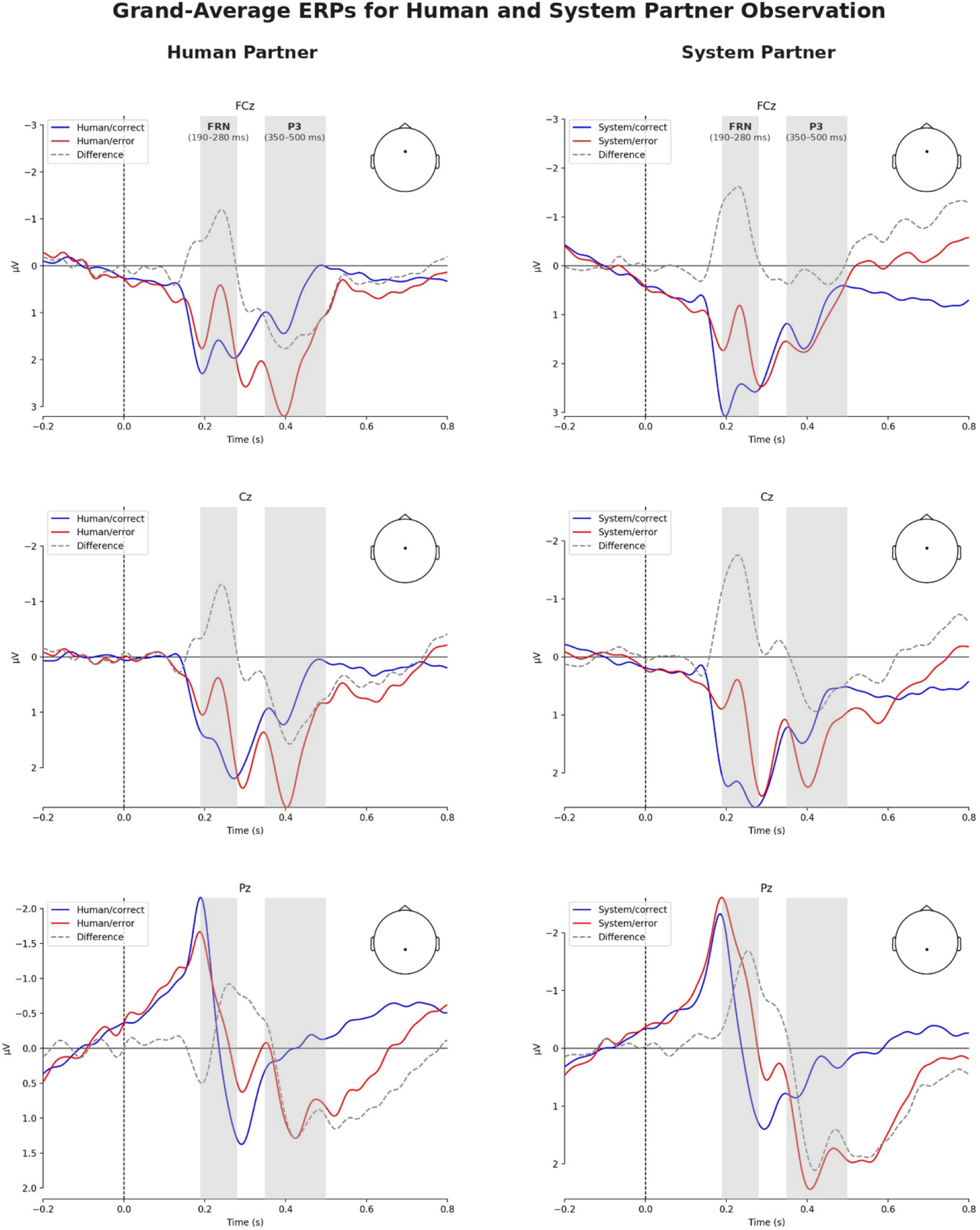
Grand average ERPs and difference waves for Human (left), and System (right), observation at electrodes FCz, Cz and Pz. A feedback-related negativity (FRN) is visible for both partner types at electrode FCz, peaking at approximately 235ms post-feedback. A P3 component is also visible for both partners with differing topography. For the Human partner a broad frontal-to-posterior P3 peaking at approximately 400ms is visible, consistent with both frontal P3a and posterior P3b sub-components. For the System partner, the frontal P3a is absent, with only the posterior P3b component visible, also peaking at approximately 400ms. Grey shaded areas indicate the analysis windows post-feedback for the FRN (190-280ms) and the P3 (350-500ms).

Topographical plots were produced in the time windows 190-280ms for the FRN and 350-500ms for the P3 based on visual inspection of the grand average ERPs. The plots, shown in Figure 5, highlight the difference for the error condition minus the correct condition by group. The FRN time window plots show a broad-based fronto-central distribution that is similar for both the human and system groups. The P3 plots however do not show a similar pattern by group, with a broad frontal-to-posterior distribution for the human group, while the system group shows a posterior distribution centred over electrode Pz.

**Figure 5.**
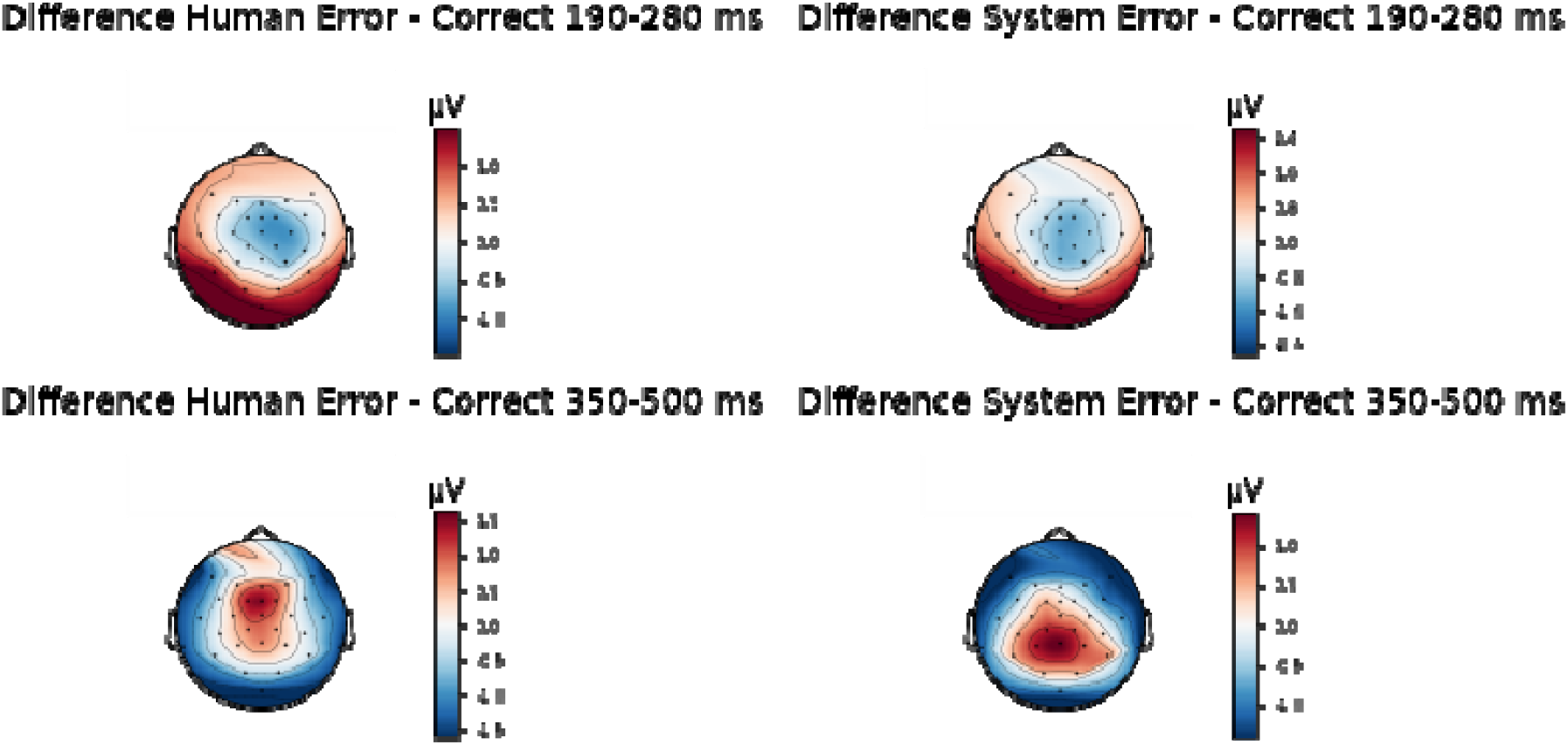
Topographical error minus correct heatmaps for Human (left), and System (right), observation during the FRN and P3 time windows. A frontal-central negativity for both the Human and System partner types is clearly visible in the 190-280ms time-window, consistent with a feedback-related-negativity (FRN) effect. Note however the differing topography in the P3 time window (350-500ms), where the Human partner shows a broad frontal-to-posterior positivity reflecting both P3a and P3b sub-components, while for the System partner the positivity has a clear P3b-like posterior distribution with no frontal positivity present.

#### FRN Time Window (190-280ms)

Modelling of the FRN time window data revealed significant main effects of accuracy [type II Wald test: *χ*^2^ (1) = 69.30, *p* < 0.001], error rate [type II Wald test: *χ*^2^ (2) = 56.40, *p* < 0.001], and sagittality [type II Wald test: *χ*^2^ (1) = 11464.90, *p* < 0.001], as well as several significant interactions (see Supplementary Table A5). No significant main effect of group or performance prediction was seen. The significant interaction of group x accuracy x error rate x pre-block prediction x sagittality [type II Wald test: *χ*^2^ (4) = 37.80, *p* < 0.001] is reported here and is visualised separately for the system and human groups in Figures 6A and 6B, respectively. Data represents fitted model values and 83% confidence intervals (see Supplementary Table A6 for the full model summary). For the system group, a broad based FRN effect can be seen, with more negative-going ERPs for error versus correct trials from frontal to posterior sites across all error rates and levels of pre-block prediction. A notable interaction of error rate and pre-block prediction can be seen over frontal sites with high error rates eliciting a larger FRN effect for better predictions than low and medium error rates. A reversal of this trend can also be seen over frontal sites with worse predictions resulting in an increased FRN effect for low error rates compared to medium and high error rates.

**Figure 6.**
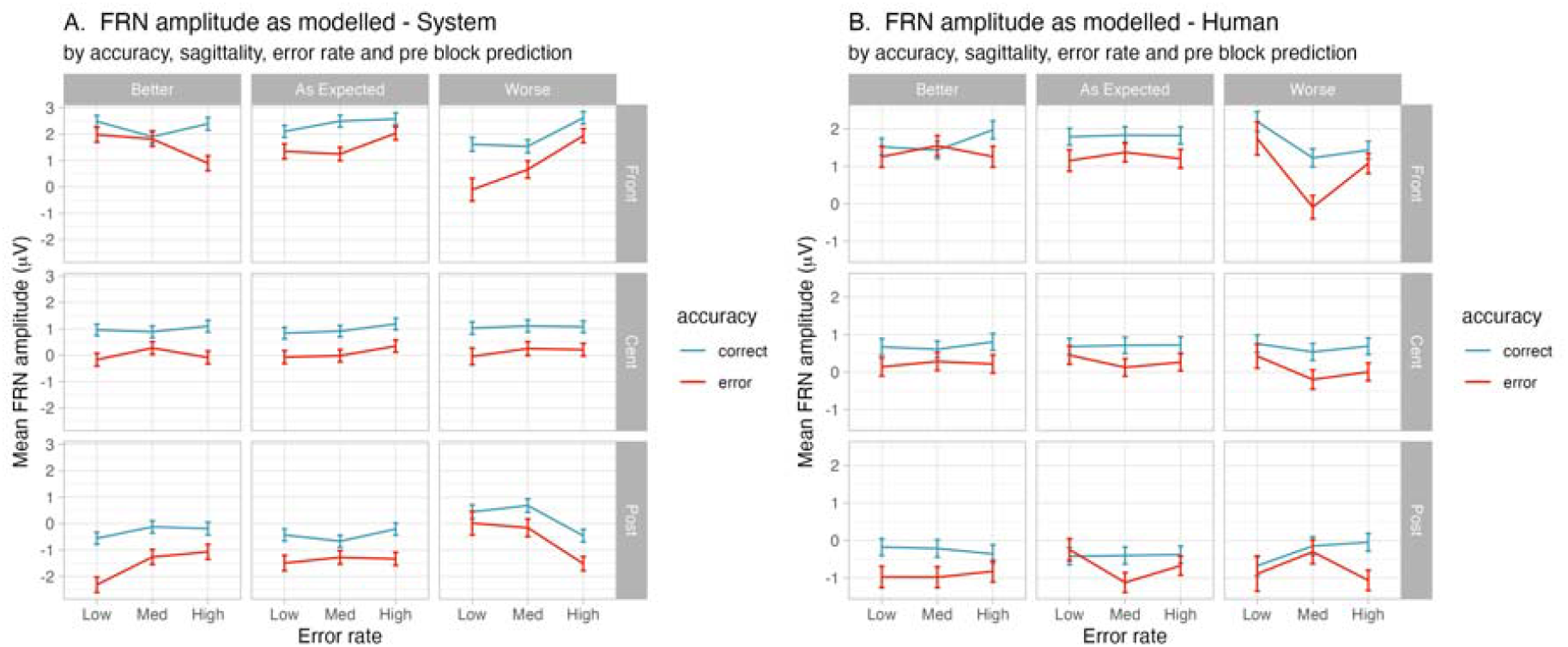
Model summaries of the FRN time window interactions by accuracy, sagittality, error rate, and pre-block prediction for the system (A), and human (B), partner groups

A similarly broad based though less consistent FRN effect is also visible for the human group with a small number of instances of error trials being less negative than correct trials. An interaction over frontal sites similar to that seen for the system group again shows low error rates eliciting an increased FRN effect compared to medium and high error rates for better predicted performance. The pattern of activity is somewhat different however for worse predictions where medium error rates show the largest FRN effect overall.

#### P3 Time Window (350-500ms)

In the P3 time window, the linear mixed-effects model also revealed significant main effects of accuracy [type II Wald test: *χ*^2^(1) = 22.30, *p* < 0.001], error rate [type II Wald test: *χ*^2^(2) = 98.60, *p* < 0.001], and sagittality [type II Wald test: *χ*^2^(1) = 184.00, *p* < 0.001] along with several significant interactions (summarised in Supplementary Table A7). Again, there were no significant main effects of group or performance prediction. The same significant interaction of group x accuracy x error rate x pre-block prediction x sagittality [type II Wald test: *χ*^2^(4) = 35.30, *p* < 0.001] seen for the FRN data is seen for the P3 data and has been visualised separately for the two groups. Data represents fitted model values and 83% confidence intervals (see Supplementary Table A8 for the full model summary). Figure 7A shows the data for the system group where a P3 effect is clearly visible over posterior sites which is largely diminished over central sites and disappears completely over frontal sites where correct trials have more positive amplitudes than errors. The interaction of error rate and pre-block prediction is minimal for both better and as expected predictions. The largest P3 effect can be seen for worse prediction and low error rates, with the effect reducing as error rates increase.

**Figure 7.**
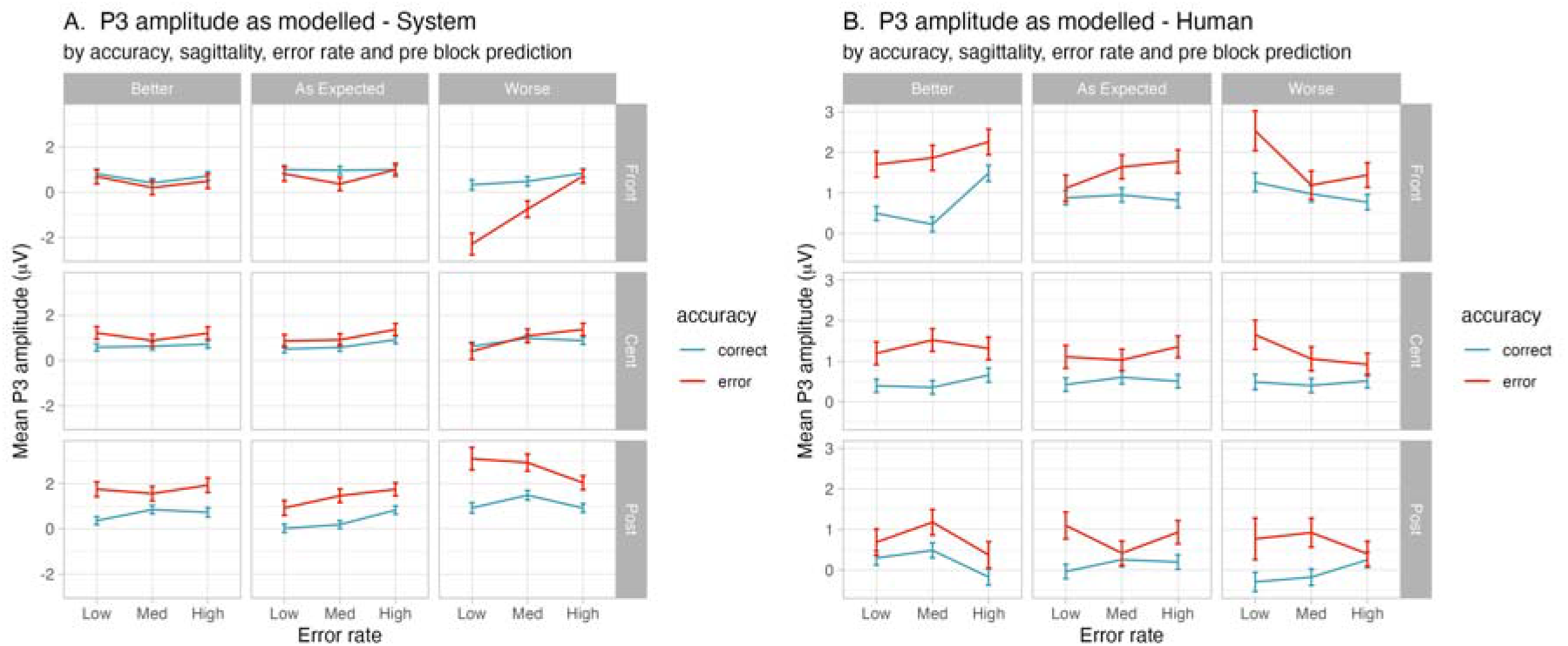
Model summaries of the P3 time window interactions by accuracy, sagittality, error rate, and pre-block prediction for the system (A), and human (B), partner groups

In contrast to the largely posterior distribution of the system group P3 activity, the human group activity visualised in Figure 7B shows a broadly distributed P3 across all sites, predictions and error rates. Worse predictions here also elicit larger P3 effects from low error rates compared to medium and high rates, however unlike the system group the activity here is widespread from frontal to posterior sites. A P3 effect is noticeable for better performance predictions to medium error rates compared to low and high error rates over frontal and central sites. This trend is reversed for as expected predictions where medium error rates result in reduced P3 effects compared to low and high rates across central and posterior sites.

## Discussion

### Summary of the main findings

In the present study, we investigated the FRN and P3 components as online measures of TiA by asking participants to observe and rate the performance of either a human or an autonomous system partner on a complex real-world image categorisation task. Behavioural results showed that, as expected, the observed error rate of a partner’s performance was a significant predictor of a participant’s pre-block performance predictions, which were worst when observed errors were increased and improved as observed errors reduced. A significant effect of error rate was also observed for the monitoring accuracy data, with more accurate monitoring of partner performance occurring when the error rates were either better, or worse, than the as expected error rate. No other effects on the behavioural data were found. Crucially, there was no behavioural effect of partner type for either the performance predictions or the monitoring accuracy. As predicted, ERP results showed a significant FRN effect and P3 effect for both the human and the autonomous system groups for observed partner errors compared to correct partner choices. While it was predicted that P3 amplitudes would reveal an increased P3 effect for human errors compared to autonomous system errors, the P3 results revealed a complex pattern of activity with distinct topographical patterns for the two groups. The human group data revealed a widespread P3 effect with errors eliciting increased amplitudes, however the effect seen for the system group was markedly different as the P3 effect was most visible over posterior sites and was absent over frontal electrodes. For both the FRN and the P3 data, complex interactions revealed interesting effects of group, error rate, and performance prediction that showed differences in observed ERP amplitudes.

### Task and behavioural effects

The behavioural data for our study show that we have been able to extend the investigation of TiA to a novel naturalistic complex real-world task. While most previous studies used a form of passive observation of a partner’s performance and variations of the Eriksen Flanker task (de Visser et al., 2018; Somon et al., 2019), we used a task that required participants to actively search for and fixate on the correct target prior to feedback of their partner’s performance. In addition to the active search required, the use of real images of cats and dogs and unique image stimuli compared to the traditional Flanker arrows adds substantial difficulty to the visual processing required by our task. Despite these changes to the task design, the monitoring accuracy data showed that participants were able to monitor and successfully differentiate the observed performance of their partner with our increased task demands. A significant effect of error rate reflected that accuracy was relatively poor for the medium error rate blocks but increased considerably for the low and high error rate blocks. This demonstrates that participants were able to identify when performance fluctuated away from the expected approximately 20% medium error rate. This result is encouraging as, in addition to our stimuli being different to previous tasks, our task implemented a markedly different method of error rate manipulation compared to previous studies. de Visser et al. (2018) used fixed 60% or 90% error rates in their task design and found that participants quickly identified the true performance level of the observed algorithm, while Somon et al. (2019) similarly used a predictable fixed rate of 33.3%. This fixed error rate approach does not allow for the investigation of a dose-response relationship between accuracy and error salience. By contrast, our non-standardised block lengths (47 to 53 trials per block), the different number of errors for each block (5 to 16), and the counterbalanced ordering of the error rates addressed this problem and potentially allowed us to more effectively manipulate the reliability, and therefore the trustworthiness, of the observed partner in our task.

The results for the pre-block predicted performance ratings indicate that our manipulation of the reliability and trustworthiness of the partner performance was relatively successful. Predicted performance rating decisions were significantly related to the observed partner error rates, suggesting that our participants were able to effectively calibrate their trust in their partner based on the observed error rates. This alignment between participants’ trust judgments and the system’s actual performance is consistent with the concept of calibrated trust as outlined by Lee and See (2004). Interpreting these results as participants having calibrated their trust in their partner suggests that our task performed as intended and is thus appropriate for an investigation of TiA. The result is also consistent with the finding of de Visser et al. (2018), who reported that subjective ratings of trustworthiness of computer algorithms were significantly influenced by the observed error rate. We hesitate to suggest however that the observed error rate was the only factor influencing participants performance predictions, as although the pattern of the data follows the direction we would expect, with increasing error rates resulting in worse predicted performance, there is some bias in participant responses evident. Mean ratings of the high error rate blocks for example barely rate below the mid-point on the prediction rating scale showing that while participants predicted worse performance for these blocks than the blocks with low or medium error rates, they still were predicting performance to be nearly at the ‘as expected’ level. A possible explanation for this bias may be that participants were actually convinced by our cover story and so kept expecting performance to return to the 20% correct ‘as expected’ performance level that they were told the partner would perform at.

The most interesting result of the behavioural data was the failure to find a significant effect of partner type for either the monitoring accuracy or the predicted performance metrics. The results show that for each group participants were equally well able to track and rate their partner’s observed performance, and they also made similar performance predictions for each group. The failure to find a significant effect of partner type in the behavioural data is not necessarily a surprise, as the randomisation of error rates and the stimuli used for the human and autonomous system groups were identical. These findings are interesting though because they do stand in stark contrast to the EEG data discussed below, where partner type was shown to significantly influence the results.

### Electrophysiological correlates of performance monitoring

The observation that FRN and P3 amplitudes were increased to partner error feedback compared to correct feedback was an expected result due to the apparent reliability of finding the FRN and P3 to visually presented feedback disambiguating outcomes. By finding this result using our image categorisation task and observation of partner performance we have added to the diverse range of tasks previously reporting an FRN and P3 (or similar error monitoring components) such as human-machine interaction (Ferrez & Millán, 2008), Flanker (de Visser et al., 2018 ; Somon et al., 2019), observed Flanker (van Schie et al., 2004) and coin-toss (Long et al., 2012), amongst others. As such this study provides further evidence that the FRN and P3 components are robust enough to be successfully elicited by our complex real-world task, giving an initial indication that these components may be suitable for use in real-world settings with practical applications. This is an important step towards more ecologically valid research into the study of TiA, however, a more nuanced exploration of the data found that both the FRN and P3 were also influenced by partner type, error rate, and predicted performance.

### FRN and P3 interactions by partner, error rate, performance prediction, and sagittality

The results of this study showed an FRN effect for both human and autonomous system partners, larger for error than correct trials over frontal electrodes, which was further modulated by performance predictions (trust) and observed performance (error rate). This result adds to the previous studies of trust in humans and TiA by directly comparing ERP responses to the two partner types in the one study. Our finding of an FRN and P3 that was modulated by trust (predicted performance) and observed performance for both partner types is in contrast to the previous work by Long et al. (2012) that found only an FRN for rewarded trust without finding a corresponding P3 component in a human partner and the study by de Visser et al. (2018) which reported only the P3 (oPe) component to be modulated by participant trust in an automated algorithm. Our results indicate that previous suggestions for the application of human correlates of performance monitoring to the study of TiA (Drnec et al., 2016; Somon et al., 2017) could be successfully applied in real-world scenarios, with the caveat that brain responses when observing humans and autonomous systems differ, meaning the direct application of findings for human performance monitoring may need adaptation for monitoring autonomous systems depending on the task at hand.

Indeed, in the case of our study we found significant interactions of the FRN and the P3 for both partner types, but the predicted performance and error rate conditions under which the differences for each partner were observed, as well as the topography of the P3 component for each partner type, differed. The FRN effect observed for the autonomous system condition was most pronounced over frontal electrodes when there was an extreme mismatch between predicted and observed performance, specifically when predicted performance was better or worse than expected and the observed performance was the opposite to predictions. This finding is consistent with FRN literature that describes enhanced amplitudes to unexpected outcomes and that the FRN is known to be scaled by the reward prediction error which can be thought of as the difference between predicted and actual outcomes (Ullsperger et al., 2014). The FRN observed in the human partner data showed similar mismatch effects, but the pattern here was slightly different. There was a noticeable effect over frontal electrodes when participants predicted to see better than expected performance and the outcome was the opposite, however the largest effect was seen when predicting worse performance and the observed performance was at the ‘as expected’ level, which indicated the human partner was actually performing as they should be. Interpreting this result in terms of the FRN being increased to unexpected outcomes, we can suggest that for the human partner once performance was seen to have been worse than it was expected to be, a return to the as expected performance level was not something that participants expected to observe.

### Topographical shift in the P3 component may reveal distinct neural processing of observed partner type

Our study yielded one particularly intriguing finding in the ERP data: a clear difference in the topography of the P3 component between human and autonomous system partners. The P3 observed for the human partner consisted of a broad topographical distribution from frontal to posterior electrodes. We interpret this as reflecting the presence of both a P3a and a P3b sub-component of the P3. By contrast, for the autonomous system partner, there was no P3a component visible and the P3 topography was limited to the central-posterior regions, reflecting only a P3b. The absence of the P3a for the autonomous system condition was unexpected. Although prior work has shown that the P3a may be absent when the FRN itself is not elicited (e.g., during fictive outcomes lacking action-based value updating; Fischer & Ullsperger, 2013), the performance monitoring literature generally indicates that a frontal P3a reliably follows an FRN (Ullsperger et al., 2014). We also expected to find a frontal P3 component based on the results of previous studies of TiA (de Visser et al., 2018; Somon et al., 2019). In the following, we suggest that this might have resulted from inherent differences in our task design compared to previous studies of TiA finding a frontal P3.

In the performance monitoring literature, the P3a following the FRN has been linked to the allocation of attentional resources to unexpected outcomes and the initiation of behavioural adjustments (Ullsperger, Danielmeier, & Jocham, 2014). Further, P3a amplitude is modulated by the salience of monitored events and is influenced by factors such as motivational and emotional significance (Nieuwenhuis, Aston-Jones, & Cohen, 2005; Polich, 2007). Previous work comparing human and system monitoring can provide informative context for understanding when frontal P3 components are observed. Somon et al. (2019) found a frontal positivity to both human and autonomous system errors, though the component was reduced for the autonomous system condition. Their task required participants to make trial-by-trial judgements of partner performance accuracy, keeping them actively engaged as the observed feedback was directly tied to required participant responses. In this way, their task ensured that frontal attentional resources were directed to the trial outcome, which maintained response readiness and, likely, generated the P3a component. The short duration (10ms) of the initial stimulus presentation used by Somon may also, at times, have induced uncertainty about the correct partner response due to participants’ difficulty discriminating the target. A similar result for autonomous system supervision was reported by de Visser et al. (2018) who found a frontal P3a-like component using a task requiring continuous monitoring of outcomes and overt evaluative responses. The authors stated that evaluative responses were required to maintain participant engagement in the monitoring task, and as such the maintenance of frontal attention allocation to the task was a deliberate design choice. A recent study by Roeder and colleagues (2025) further clarifies conditions under which frontal P3a activity may be observed in a human-AI interaction task. Their study reported a sequence of feedback-related components they termed an N2-P3a-P3b complex (though following our parsimony account, we would consider the N2 to be an FRN) when participants’ own stimulus classifications mismatched the AI-generated outputs. Notably, participants in that paradigm were not required to provide overt responses at the time of feedback. The task required passive observation of the AI’s classification outcome, however, because participants had already generated a classification decision on each trial, incongruent AI feedback necessarily conflicted with self-generated expectations, requiring evaluation and updating of one’s own belief state. Further, the inherent difficulty of judging real faces from AI-generated faces possibly caused genuine uncertainty about whose classification was indeed correct, the participants’ or the AI’s. The resulting P3a may have been driven by conflict monitoring and expectancy violations resulting from self–other disagreement and decision uncertainty.

The results of these previous studies indicate that in paradigms requiring overt behavioural responses to errors or involving feedback that conflicts with self-generated expectations, the P3a serves both attentional allocation and action-updating or conflict-resolution functions. This makes it difficult to isolate whether social or motivational factors independently modulate frontal engagement during pure passive observation of another agent’s performance. By contrast, our passive monitoring task required no behavioural adjustments from participants in either condition. It also involved no self-generated classification or expectations that could conflict with observed outcomes. In our task, the outcome was always known by participants as the classification task was simple for them to perform accurately themselves, leaving no doubt as to the accuracy of the observed partner performance. By eliminating both action demands and self-generated conflict our task may have permitted isolation of the pure attentional allocation component of the P3a during observation of another agent’s performance. As the P3a was preserved for the human condition, despite the lack of action requirements and absence of self-generated conflict, this suggests that social and motivational salience alone, independent of behavioural adjustment needs or expectancy violation, is sufficient to maintain frontal attentional engagement. It is possible the human partner errors may have been salient due to factors including perceived intentionality and agency, a need for social comparison and self-evaluation (Valt, Sprengel, & Sturmer, 2020), and the engagement of empathy and social cognition.

In stark contrast, despite equivalent behavioural trust calibration and intact FRN responses, the P3a was absent when monitoring autonomous systems. This contrasts with the reduced but present P3a observed when system monitoring includes action demands (Somon et al., 2019; de Visser et al., 2018) or self-generated conflict (Roeder et al., preprint). Our findings suggest that when action requirements, self-generated conflict, and social salience are all absent, there may be no mechanism remaining during passive observation of system performance to sustain frontal attentional engagement. Further, our findings qualify the perspective that error-related potentials could serve as neural indices of TiA (Drnec et al., 2016; Somon et al., 2017).

This perspective, consistent with CASA theory’s prediction of social equivalence in human-computer interaction (Nass & Moon, 2000), suggests that known error-monitoring components should transfer equivalently when monitoring human versus system partners. While we have shown that components such as the FRN and P3 are potentially useful indicators of TiA, providing support for their proposed adoption to the field of TiA measurement, the absence of the P3a component when monitoring autonomous systems in our results indicates human performance monitoring mechanisms may not equivalently transfer to the monitoring of an automated system as suggested. Instead, it appears that different neural mechanisms are engaged when monitoring a human versus an autonomous system. This finding suggests boundary conditions exist for the adaptation of neuroscience methods to TiA and also potentially indicates limited generalisability of CASA theory at a neural level in this context.

A possible explanation for the observed divergence in P3a activity could be that social and attentional processes are differentially engaged when monitoring a human compared to an automated system. When monitoring human partners, it is likely that social cognition processes such as theory of mind (Van Overwalle & Baetens, 2009), empathy (Singer & Lamm, 2009), and social evaluation (Ruff & Fehr, 2014) may maintain active attentional engagement as reflected in the preserved P3a we observed here for human partners. This interpretation is consistent with findings that error-related potentials are modulated by social context (de Bruijn, Schubotz, & Ullsperger, 2007), empathy (Rak, Bellebaum, & Thoma, 2013), as well as the social consequences of observed errors (Koban, Pourtois, Vocat, & Vuilleumier, 2010). When monitoring autonomous systems, it is possible these social and emotional processes are not engaged, or engaged to a lesser degree, meaning the psychological mechanisms facilitating active monitoring may be absent. This could reduce monitoring engagement, enabling a condition whereby active attentional resources are minimised rather than maintained. Critically, our results indicate this reduction in engagement is selective as basic error detection (FRN) and posterior evaluation (P3b) remain intact, but frontal attentional engagement (P3a) is absent. This pattern suggests that some neural mechanisms of human performance monitoring do adapt to system monitoring (FRN, P3b), whereas others may require social and emotional engagement to be maintained (P3a).

The absent P3a seen here may therefore be a potential neural marker of automation complacency (Parasuraman & Riley, 1997; Parasuraman, Sheridan, & Wickens, 2000), whereby humans fail to maintain adequate vigilance when monitoring automation. Our findings suggest this phenomenon may not reflect merely motivational failure as previously theorised, but also a fundamental shift in cognitive processing mode that occurs specifically when human partners are replaced by technological systems (cf. differential neural processing of in-group vs. out-group members; Amodio, 2014). Understanding which neural mechanisms may transfer to the monitoring of automation and which require social engagement has critical implications for developing neural indices of TiA, and for designing effective human-automation teams. Caution should be used when assuming complete equivalence between human and system monitoring as this may overlook the potential absence of frontal P3 components during certain task scenarios, which may be a neural signature of automation complacency itself.

This pattern of selectively reduced frontal engagement when monitoring technological versus human agents may reflect a tendency to minimise cognitive effort when interacting with technological systems. Such a pattern of technology-related cognitive economisation (TRCE) conceptually aligns with established principles of human cognition. Humans are cognitive misers who seek to minimise mental effort, when possible (Shenhav et al., 2017), and effort avoidance is a fundamental feature of cognitive processing such that people naturally default to less effortful cognitive strategies when opportunity permits (Kool & Botvinick, 2014; Shah & Oppenheimer, 2008). The absent P3a observed in the present study fits this framework as when social and emotional processes that normally sustain effortful engagement with human partners are not engaged, cognitive effort minimisation may be permitted. This potentially manifests as reduced frontal attentional investment during system monitoring. While our findings potentially demonstrate TRCE in a passive monitoring context, this phenomenon may have broader relevance for understanding human-technology interaction. It is possible that TRCE may extend beyond passive monitoring to encompass other forms of effort minimisation with technological systems, such as cognitive offloading whereby people delegate cognitive work to AI assistants or large language models (Risko & Gilbert, 2016), or uncritical acceptance of algorithmically-generated content without adequate verification (Vasconcelos et al., 2022). Both passive monitoring disengagement (as demonstrated here) and deliberate cognitive delegation through offloading to AI systems, could reflect the same underlying tendency towards effort economisation when interacting with technology.

We note several important caveats to the above. It is likely that the extent of TRCE may be moderated by design features such as anthropomorphic characteristics or human-like conversational language that engage social and cognitive processing (Epley, Waytz, & Cacioppo, 2007). Systems designed with such features might recruit the social and emotional networks that maintain cognitive engagement, potentially reducing TRCE. Additionally, the present study was limited to examining passive monitoring of partner performance rather than considering active trust decision-making in collaborative contexts. It is thus unknown whether TRCE would extend from passive observation to an active evaluation paradigm requiring judgments of system trustworthiness or reliance. Future research should examine TRCE in different engagement contexts, including active decision-making during human-automation teaming scenarios, which could establish the boundary conditions and generalisability of the pattern found here. Furthering this finding to understand when and how humans economise cognitive resources with technology has critical implications for automation safety, human-AI collaboration, and the design of systems that maintain appropriate human oversight.

### Practical implications/future directions

The results of the present study suggest that EEG-based measures can potentially be used as sensitive indicators of TiA. The finding of an effect of partner type in our FRN and P3 data that was modulated by participant trust in their partner and observed error rate, without finding behavioural differences for these effects, is a particularly promising finding for future investigations of TiA as it highlights the potential added benefit of EEG-based measures over traditional methods. Further, the effectiveness of our novel task design, which reliably elicited the desired behavioural effects and targeted ERPs, suggests that future studies may be able to successfully use similarly complex tasks to investigate other types of human-autonomous system interaction. This is encouraging, as continued research in this domain may allow for the future development of diagnostic tools that offer real-time measurement of TiA, which are complementary to behavioural measures, while not being limited by the known issues inherent to trust measurement (e.g. bias in participant responses, post hoc measurement).

The novel finding that the P3a was absent for the system group has several practical implications. The absent P3a may be a marker of automation complacency that reflects a basic shift in processing during monitoring of automated or technological agents compared to humans. Understanding how, when, and under what circumstances this shift in processing mode manifests can directly inform the design of safe human-automation teams. Following the TRCE framework outlined above it appears that basic error detection and evaluative processes transfer from human monitoring to system monitoring while frontal attention does not, possibly due to reduced social engagement processes. For the future design of safe human-automation teams it may be necessary to create conditions that sustain social and cognitive engagement with a given system or technology, with the use of anthropomorphic or human-like design characteristics, such as the use of natural language, being possible candidates to achieve sustained frontal attentional engagement. Employing differing implementations of these characteristics may then mitigate TRCE, for while often considered to be aesthetic or communicative features of design these may be determinative of the vigilance a system elicits from the human user. This is an area of human-automation team design that is currently under-explored.

Future research is needed to replicate our absent P3a finding under different task and system design scenarios so that it can be established whether the absence of the P3a is a robust neural marker of reduced frontal attention during system monitoring before its use as a practical indicator of automation complacency. Beyond replication of our finding, an open question is whether the observed TRCE we reported here as reduced frontal attention and the absent P3a can be extended from this passive monitoring context to scenarios requiring active decision-making such as overt decisions to trust a system partner during collaborative teaming tasks. It is possible that in such scenarios requiring explicit responses or interventions from a human teammate or partner that the P3a may be recovered, which could facilitate the maintenance of appropriate human oversight in safety critical environments through sustained frontal attention. Research on the use of anthropomorphic or human-like characteristics in system design is also needed to establish a baseline of neuroscientific knowledge that along with specific task demands or teaming scenarios can act as moderators of TRCE. This baseline of knowledge would allow the development and design of systems that maintain appropriate levels of human oversight. Taking a meticulous approach to establishing a sound neuroscientific foundation for this emerging field is important considering the rapid pace of development and prevalence of autonomous systems and artificial intelligence in our society, particularly in safety-critical domains.

## Conclusions

The present study has made valuable contributions to the study of TiA. We have confirmed that beyond the traditional flanker task used by previous studies it is possible to obtain FRN and P3 effects from observed performance of an autonomous system during a complex task approximating a real-world human-autonomous system teaming environment. Our participants appropriately calibrated their trust in their partner based on the observed performance, and the FRN and P3 activity revealed distinct neural patterns for each partner type, modulated by error rate and predicted performance, that were not identified by the behavioural data alone. This finding demonstrates the potential for EEG to be used as a real-time, sensitive measure of TiA in complex tasks without the limitations of traditional self-report methods. Further, the novel finding that the P3a was absent following the FRN for the autonomous system group is theoretically significant as it suggests fundamental differences exist in the processing of observed human and autonomous system errors. We theorised that a selective absence of the P3a, co-occurring with FRN and P3b responses indicating retained basic error detection and outcome evaluation processes, suggests it is specifically frontal attention as indexed by the P3a that is not sustained during passive monitoring of a system partner. We suggest this may reflect Technology-Related Cognitive Economisation (TRCE), a tendency to default to a reduced state of cognitive effort which manifests when the social and emotional processes that sustain effortful engagement with humans are not present to similarly sustain engagement in the context of system monitoring. This neural pattern of effort economisation may be a marker of automation complacency that also potentially identifies boundary conditions under which the application of human performance monitoring theories may not apply to system supervision. Understanding how this pattern of cognitive economisation occurs and investigating the factors that may mitigate it, such as system characteristics and task design, has potentially important implications for safety in human automation teams and the development of neural measures of TiA.

