## Supplementary materials for "The absent P3a. Performance monitoring ERPs differentiate trust in humans and autonomous systems"

2025-06-11

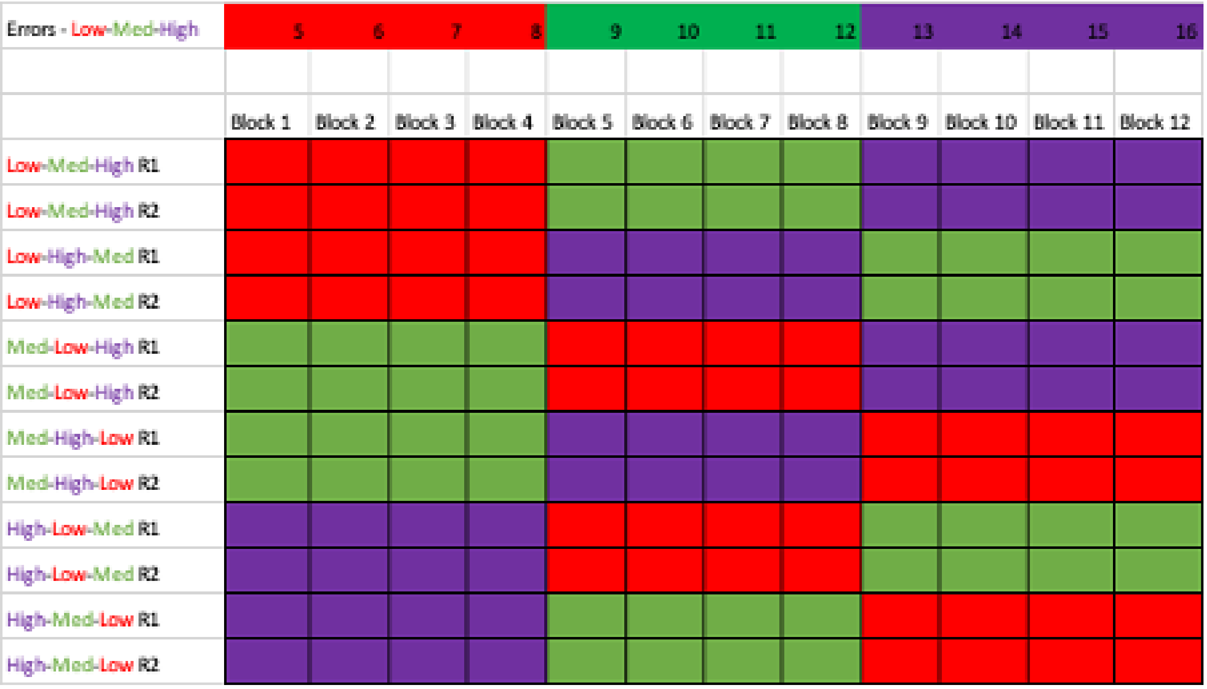


Figure S1: Partner error rate block orders

Table S1: Analysis of Deviance Table (Type II Wald chisquare tests)Response: Monitoring Accuracy

| | $\chi^2$ | df | Pr(> $\chi^2$) |
|---|---|---|---|
| group | 0.034 | 1 | 0.85000 |
| rate | 17.200 | 2 | 0.00018 |
| group:rate | 3.100 | 2 | 0.21000 |

Table S2: Summary of Monitoring Accuracy model produced by the call `glmer(formula = Monitor_Accuracy ~ group * rate + (1 | id), data = logfile_behav, family = "binomial")`
Generalized linear mixed model fit by maximum likelihood (Laplace Approximation)

| AIC | BIC | logLik | deviance | df.resid |
|---|---|---|---|---|
| 771 | 801 | -378 | 757 | 569 |

Scaled residuals:

| Min | 1Q | Median | 3Q | Max |
|---|---|---|---|---|
| -1.9 | -1.05 | 0.58 | 0.78 | 1.54 |

Random effects:

| Groups | Term | Std.Dev. |
|---|---|---|
| id | (Intercept) | 0.52487 |

Number of obs: 576, groups: id, 48.

Fixed effects:

| | Estimate | Std. Error | z value | Pr(>\|z\|) | |
|---|---|---|---|---|---|
| (Intercept) | 0.79 | 0.25 | 3.2 | 0.0016 | ** |
| groupsystem | −0.3 | 0.35 | −0.85 | 0.39 | |
| rateHigh | −0.2 | 0.31 | −0.63 | 0.53 | |
| rateMed | −1.2 | 0.31 | −3.8 | 0.00013 | *** |
| groupsystem:rateHigh | 0.24 | 0.44 | 0.56 | 0.58 | |
| groupsystem:rateMed | 0.75 | 0.43 | 1.7 | 0.086 | . |

Table S3: Analysis of Deviance Table (Type II Wald chisquare tests)
Response: Pre-Block Performance Prediction

| | $\chi^2$ | df | $Pr(>\chi^2)$ |
|---|---|---|---|
| group | 0.0084 | 1 | 0.93 |
| rate | 41.9000 | 2 | 0.00 |
| group:rate | 0.2000 | 2 | 0.91 |

Table S4: Summary of Pre-block Performance Prediction model produced by the call `lmer(formula = Pre_Block_Pred ~ group * rate + (1 | id), data = logfile_behav, REML = FALSE, control = lmerControl(optimizer = "bobyqa", calc.derivs = TRUE))`
Linear mixed model fit by maximum likelihood . t-tests use Satterthwaite's method

| AIC | BIC | logLik | deviance | df.resid |
|---|---|---|---|---|
| 1174 | 1209 | -579 | 1158 | 565 |

Scaled residuals:

| Min | 1Q | Median | 3Q | Max |
|---|---|---|---|---|
| -2.29 | -0.61 | -0.01 | 0.82 | 1.95 |

Random effects:

| Groups | Term | Std.Dev. |
|---|---|---|
| id | (Intercept) | 0.28032 |
| Residual | | 0.63169 |

Number of obs: 573, groups: id, 48.

Fixed effects:

| | Estimate | Std. Error | df | t value | Pr(>\|t\|) | |
|---|---|---|---|---|---|---|
| (Intercept) | 0.37 | 0.086 | 1.2e+02 | 4.4 | 2.9e−05 | *** |
| groupsystem | −0.042 | 0.12 | 1.2e+02 | −0.34 | 0.73 | |
| rateHigh | −0.44 | 0.091 | 5.3e+02 | −4.8 | 1.8e−06 | *** |
| rateMed | −0.23 | 0.091 | 5.3e+02 | −2.5 | 0.013 | * |
| groupsystem:rateHigh | 0.047 | 0.13 | 5.3e+02 | 0.36 | 0.72 | |
| groupsystem:rateMed | 0.052 | 0.13 | 5.3e+02 | 0.4 | 0.69 | |

Table S5: Analysis of Deviance Table (Type II Wald chisquare tests)Response: FRN amplitude

| | $\chi^2$ | df | $Pr(> \chi^2)$ |
|---|---|---|---|
| accuracy | 69.30 | 1 | 0.00000 |
| sag | 11464.90 | 1 | 0.00000 |
| group | 0.46 | 1 | 0.50000 |
| rate | 56.40 | 2 | 0.00000 |
| Pre__pred__factor | 2.90 | 2 | 0.24000 |
| accuracy:sag | 6.30 | 1 | 0.01200 |
| accuracy:group | 5.60 | 1 | 0.01700 |
| sag:group | 239.20 | 1 | 0.00000 |
| accuracy:rate | 1.80 | 2 | 0.40000 |
| sag:rate | 40.70 | 2 | 0.00000 |
| group:rate | 23.80 | 2 | 0.00001 |
| accuracy:Pre__pred__factor | 2.70 | 2 | 0.26000 |
| sag:Pre__pred__factor | 127.20 | 2 | 0.00000 |
| group:Pre__pred__factor | 11.20 | 2 | 0.00360 |
| rate:Pre__pred__factor | 12.40 | 4 | 0.01500 |
| accuracy:sag:group | 4.00 | 1 | 0.04500 |
| accuracy:sag:rate | 0.38 | 2 | 0.83000 |
| accuracy:group:rate | 9.90 | 2 | 0.00720 |
| sag:group:rate | 19.60 | 2 | 0.00006 |
| accuracy:sag:Pre__pred__factor | 23.60 | 2 | 0.00001 |
| accuracy:group:Pre__pred__factor | 3.10 | 2 | 0.21000 |
| sag:group:Pre__pred__factor | 18.70 | 2 | 0.00009 |
| accuracy:rate:Pre__pred__factor | 34.30 | 4 | 0.00000 |
| sag:rate:Pre__pred__factor | 167.20 | 4 | 0.00000 |
| group:rate:Pre__pred__factor | 32.30 | 4 | 0.00000 |
| accuracy:sag:group:rate | 9.90 | 2 | 0.00720 |
| accuracy:sag:group:Pre__pred__factor | 0.98 | 2 | 0.61000 |
| accuracy:sag:rate:Pre__pred__factor | 72.90 | 4 | 0.00000 |
| accuracy:group:rate:Pre__pred__factor | 4.30 | 4 | 0.36000 |
| sag:group:rate:Pre__pred__factor | 209.50 | 4 | 0.00000 |
| accuracy:sag:group:rate:Pre__pred__factor | 37.80 | 4 | 0.00000 |

Table S6: Summary of FRN time window model produced by the call `lmer(formula = eeg ~ accuracy * sag * group * rate * Pre_pred_factor + (1 + accuracy | id), data = subset(maxeeg1, wname == "FRN"), REML = FALSE, control = lmerControl(optimizer = "bobyqa", calc.derivs = TRUE))`
Linear mixed model fit by maximum likelihood . t-tests use Satterthwaite's method

| AIC | BIC | logLik | deviance | df.resid |
|---|---|---|---|---|
| 2356146 | 2356973 | -1177997 | 2355994 | 391690 |

Scaled residuals:

| Min | 1Q | Median | 3Q | Max |
|---|---|---|---|---|
| -4.58 | -0.61 | 0 | 0.61 | 4.39 |

Random effects:

| Groups | Term | Std.Dev. | Corr |
|---|---|---|---|
| id | (Intercept) | 0.76553 | |
| | accuracyerror | 0.57141 | -0.360 |
| Residual | | 4.89127 | |

Number of obs: 391766, groups: id, 48.

Fixed effects:

| | Estimate | Std. Error | df | t value | Pr(>\|t\|) | |
|---|---|---|---|---|---|---|
| (Intercept) | 0.71 | 0.16 | 51 | 4.5 | 4.2e−05 | *** |
| accuracyerror | −0.59 | 0.14 | 80 | −4.3 | 4.6e−05 | *** |
| sag | 1.6 | 0.062 | 3.9e+05 | 26 | 1.2e−143 | *** |
| groupsystem | 0.2 | 0.23 | 51 | 0.88 | 0.38 | |
| rateLow | −0.032 | 0.046 | 3.9e+05 | −0.69 | 0.49 | |
| rateHigh | 0.0071 | 0.047 | 3.9e+05 | 0.15 | 0.88 | |
| PrepredfactorBetter | −0.11 | 0.052 | 3.8e+05 | −2.1 | 0.04 | * |
| PrepredfactorWorse | −0.18 | 0.063 | 3.8e+05 | −2.8 | 0.0053 | ** |
| accuracyerror:sag | 0.19 | 0.14 | 3.9e+05 | 1.4 | 0.17 | |
| accuracyerror:groupsystem | −0.34 | 0.19 | 78 | −1.8 | 0.079 | . |
| sag:groupsystem | 0.66 | 0.086 | 3.9e+05 | 7.7 | 1.3e−14 | *** |
| accuracyerror:rateLow | 0.36 | 0.11 | 3.6e+05 | 3.2 | 0.0016 | ** |
| accuracyerror:rateHigh | 0.13 | 0.095 | 3.7e+05 | 1.4 | 0.17 | |
| sag:rateLow | −0.017 | 0.088 | 3.9e+05 | −0.19 | 0.85 | |
| sag:rateHigh | −0.022 | 0.091 | 3.9e+05 | −0.24 | 0.81 | |
| groupsystem:rateLow | −0.044 | 0.064 | 3.9e+05 | −0.69 | 0.49 | |
| groupsystem:rateHigh | 0.26 | 0.066 | 3.9e+05 | 4 | 5.7e−05 | *** |
| accuracyerror:PrepredfactorBetter | 0.26 | 0.11 | 2.9e+05 | 2.4 | 0.019 | * |
| accuracyerror:PrepredfactorWorse | −0.15 | 0.14 | 2.4e+05 | −1.1 | 0.28 | |
| sag:PrepredfactorBetter | −0.42 | 0.097 | 3.9e+05 | −4.3 | 1.4e−05 | *** |
| sag:PrepredfactorWorse | −0.61 | 0.12 | 3.9e+05 | −5.3 | 1.2e−07 | *** |
| groupsystem:PrepredfactorBetter | 0.083 | 0.074 | 3.8e+05 | 1.1 | 0.26 | |
| groupsystem:PrepredfactorWorse | 0.37 | 0.092 | 3.8e+05 | 4.1 | 4.7e−05 | *** |
| rateLow:PrepredfactorBetter | 0.095 | 0.069 | 3.9e+05 | 1.4 | 0.17 | |
| rateHigh:PrepredfactorBetter | 0.19 | 0.079 | 3.9e+05 | 2.4 | 0.017 | * |
| rateLow:PrepredfactorWorse | 0.25 | 0.1 | 3.8e+05 | 2.5 | 0.012 | * |
| rateHigh:PrepredfactorWorse | 0.15 | 0.084 | 3.8e+05 | 1.7 | 0.084 | . |
| accuracyerror:sag:groupsystem | −0.64 | 0.19 | 3.9e+05 | −3.4 | 0.00068 | *** |
| accuracyerror:sag:rateLow | −0.77 | 0.22 | 3.9e+05 | −3.5 | 0.00047 | *** |
| accuracyerror:sag:rateHigh | −0.42 | 0.18 | 3.9e+05 | −2.3 | 0.022 | * |
| accuracyerror:groupsystem:rateLow | −0.34 | 0.16 | 3.6e+05 | −2.1 | 0.032 | * |
| accuracyerror:groupsystem:rateHigh | −0.038 | 0.13 | 3.6e+05 | −0.29 | 0.77 | |
| sag:groupsystem:rateLow | −0.43 | 0.12 | 3.9e+05 | −3.5 | 0.00042 | *** |
| sag:groupsystem:rateHigh | −0.25 | 0.13 | 3.9e+05 | −1.9 | 0.052 | . |
| accuracyerror:sag:PrepredfactorBetter | 0.45 | 0.21 | 3.9e+05 | 2.1 | 0.034 | * |
| accuracyerror:sag:PrepredfactorWorse | −1 | 0.25 | 3.9e+05 | −4 | 6.1e−05 | *** |
| accuracyerror:groupsystem:PrepredfactorBetter | 0.05 | 0.16 | 2.5e+05 | 0.31 | 0.76 | |
| accuracyerror:groupsystem:PrepredfactorWorse | 0.22 | 0.2 | 2.2e+05 | 1.1 | 0.27 | |
| sag:groupsystem:PrepredfactorBetter | −0.39 | 0.14 | 3.9e+05 | −2.9 | 0.0043 | ** |
| sag:groupsystem:PrepredfactorWorse | −1 | 0.17 | 3.9e+05 | −6.1 | 8.3e−10 | *** |
| accuracyerror:rateLow:PrepredfactorBetter | −0.57 | 0.17 | 3.4e+05 | −3.4 | 0.00067 | *** |
| accuracyerror:rateHigh:PrepredfactorBetter | −0.39 | 0.16 | 3.2e+05 | −2.5 | 0.013 | * |
| accuracyerror:rateLow:PrepredfactorWorse | 0.046 | 0.25 | 3.2e+05 | 0.18 | 0.85 | |
| accuracyerror:rateHigh:PrepredfactorWorse | −0.081 | 0.17 | 2.4e+05 | −0.47 | 0.64 | |
| sag:rateLow:PrepredfactorBetter | 0.055 | 0.13 | 3.9e+05 | 0.42 | 0.67 | |
| sag:rateHigh:PrepredfactorBetter | 0.51 | 0.15 | 3.9e+05 | 3.4 | 0.00071 | *** |
| sag:rateLow:PrepredfactorWorse | 1.1 | 0.19 | 3.9e+05 | 5.8 | 6e−09 | *** |
| sag:rateHigh:PrepredfactorWorse | 0.098 | 0.16 | 3.9e+05 | 0.62 | 0.53 | |
| groupsystem:rateLow:PrepredfactorBetter | 0.054 | 0.096 | 3.9e+05 | 0.56 | 0.58 | |
| groupsystem:rateHigh:PrepredfactorBetter | −0.25 | 0.11 | 3.9e+05 | −2.2 | 0.027 | * |
| groupsystem:rateLow:PrepredfactorWorse | −0.26 | 0.14 | 3.8e+05 | −1.8 | 0.072 | . |
| groupsystem:rateHigh:PrepredfactorWorse | −0.45 | 0.12 | 3.8e+05 | −3.7 | 0.0002 | *** |
| accuracyerror:sag:groupsystem:rateLow | 1.4 | 0.31 | 3.9e+05 | 4.7 | 2.5e−06 | *** |
| accuracyerror:sag:groupsystem:rateHigh | 1.3 | 0.26 | 3.9e+05 | 5 | 4.4e−07 | *** |
| accuracyerror:sag:groupsystem:PrepredfactorBetter | 0.76 | 0.3 | 3.9e+05 | 2.6 | 0.011 | * |
| accuracyerror:sag:groupsystem:PrepredfactorWorse | 1.4 | 0.37 | 3.9e+05 | 3.9 | 8.3e−05 | *** |
| accuracyerror:sag:rateLow:PrepredfactorBetter | 0.51 | 0.32 | 3.9e+05 | 1.6 | 0.11 | |
| accuracyerror:sag:rateHigh:PrepredfactorBetter | −0.39 | 0.3 | 3.9e+05 | −1.3 | 0.2 | |
| accuracyerror:sag:rateLow:PrepredfactorWorse | 1.4 | 0.47 | 3.9e+05 | 3 | 0.0025 | ** |
| accuracyerror:sag:rateHigh:PrepredfactorWorse | 1.7 | 0.32 | 3.9e+05 | 5.4 | 8.2e−08 | *** |
| accuracyerror:groupsystem:rateLow:PrepredfactorBetter | 0.033 | 0.23 | 3.6e+05 | 0.14 | 0.89 | |
| accuracyerror:groupsystem:rateHigh:PrepredfactorBetter | −0.27 | 0.22 | 3.3e+05 | −1.2 | 0.23 | |
| accuracyerror:groupsystem:rateLow:PrepredfactorWorse | −0.28 | 0.35 | 2.8e+05 | −0.79 | 0.43 | |
| accuracyerror:groupsystem:rateHigh:PrepredfactorWorse | −0.018 | 0.25 | 2.4e+05 | −0.074 | 0.94 | |
| sag:groupsystem:rateLow:PrepredfactorBetter | 1.1 | 0.18 | 3.9e+05 | 6.1 | 8.9e−10 | *** |
| sag:groupsystem:rateHigh:PrepredfactorBetter | 0.15 | 0.21 | 3.9e+05 | 0.72 | 0.47 | |
| sag:groupsystem:rateLow:PrepredfactorWorse | −0.42 | 0.26 | 3.9e+05 | −1.6 | 0.11 | |
| sag:groupsystem:rateHigh:PrepredfactorWorse | 1.8 | 0.23 | 3.9e+05 | 7.8 | 6.1e−15 | *** |
| accuracyerror:sag:groupsystem:rateLow:PrepredfactorBetter | −1 | 0.45 | 3.9e+05 | −2.3 | 0.02 | * |
| accuracyerror:sag:groupsystem:rateHigh:PrepredfactorBetter | −1.7 | 0.43 | 3.9e+05 | −3.9 | 8.4e−05 | *** |
| accuracyerror:sag:groupsystem:rateLow:PrepredfactorWorse | −3 | 0.66 | 3.9e+05 | −4.5 | 6e−06 | *** |
| accuracyerror:sag:groupsystem:rateHigh:PrepredfactorWorse | −2.3 | 0.46 | 3.9e+05 | −5 | 6.5e−07 | *** |

Table S7: Analysis of Deviance Table (Type II Wald chisquare tests)Response: P3 amplitude

| | $\chi^2$ | df | $\Pr(>\chi^2)$ |
|---|---|---|---|
| accuracy | 22.30 | 1 | 0.00000 |
| sag | 184.00 | 1 | 0.00000 |
| group | 1.10 | 1 | 0.29000 |
| rate | 98.60 | 2 | 0.00000 |
| Pre_pred_factor | 0.35 | 2 | 0.84000 |
| accuracy:sag | 103.40 | 1 | 0.00000 |
| accuracy:group | 2.40 | 1 | 0.12000 |
| sag:group | 266.40 | 1 | 0.00000 |
| accuracy:rate | 5.30 | 2 | 0.07000 |
| sag:rate | 38.90 | 2 | 0.00000 |
| group:rate | 21.60 | 2 | 0.00002 |
| accuracy:Pre_pred_factor | 17.30 | 2 | 0.00018 |
| sag:Pre_pred_factor | 107.20 | 2 | 0.00000 |
| group:Pre_pred_factor | 13.70 | 2 | 0.00100 |
| rate:Pre_pred_factor | 15.00 | 4 | 0.00460 |
| accuracy:sag:group | 218.00 | 1 | 0.00000 |
| accuracy:sag:rate | 16.70 | 2 | 0.00024 |
| accuracy:group:rate | 6.20 | 2 | 0.04400 |
| sag:group:rate | 15.50 | 2 | 0.00042 |
| accuracy:sag:Pre_pred_factor | 26.90 | 2 | 0.00000 |
| accuracy:group:Pre_pred_factor | 2.10 | 2 | 0.35000 |
| sag:group:Pre_pred_factor | 108.50 | 2 | 0.00000 |
| accuracy:rate:Pre_pred_factor | 12.00 | 4 | 0.01700 |
| sag:rate:Pre_pred_factor | 89.10 | 4 | 0.00000 |
| group:rate:Pre_pred_factor | 43.00 | 4 | 0.00000 |
| accuracy:sag:group:rate | 6.90 | 2 | 0.03200 |
| accuracy:sag:group:Pre_pred_factor | 8.40 | 2 | 0.01500 |
| accuracy:sag:rate:Pre_pred_factor | 37.60 | 4 | 0.00000 |
| accuracy:group:rate:Pre_pred_factor | 47.70 | 4 | 0.00000 |
| sag:group:rate:Pre_pred_factor | 137.40 | 4 | 0.00000 |
| accuracy:sag:group:rate:Pre_pred_factor | 35.30 | 4 | 0.00000 |

Table S8: Summary of P3 time window model produced by the call lmer(formula = eeg ~ accuracy * sag * group * rate * Pre_pred_factor + (1 + accuracy | id), data = subset(maxeeg1, wname == "P3b"), REML = FALSE, control = lmerControl(optimizer = "bobyqa", calc.derivs = TRUE))
Linear mixed model fit by maximum likelihood . t-tests use Satterthwaite's method

| AIC | BIC | logLik | deviance | df.resid |
|---|---|---|---|---|
| 2410727 | 2411553 | -1205287 | 2410575 | 390590 |

Scaled residuals:

| Min | 1Q | Median | 3Q | Max |
|---|---|---|---|---|
| -4.1 | -0.61 | 0 | 0.6 | 4.09 |

Random effects:

| Groups | Term | Std.Dev. | Corr |
|---|---|---|---|
| id | (Intercept) | 0.55748 | |
| | accuracyerror | 0.78223 | -0.127 |
| Residual | | 5.28984 | |

Number of obs: 390666, groups: id, 48.

Fixed effects:

| | Estimate | Std. Error | df | t value | Pr(>\|t\|) | |
|---|---|---|---|---|---|---|
| (Intercept) | 0.6 | 0.12 | 56 | 5.1 | 4.8e−06 | *** |
| accuracyerror | 0.43 | 0.18 | 68 | 2.4 | 0.019 | * |
| sag | 0.49 | 0.068 | 3.9e+05 | 7.3 | 2.7e−13 | *** |
| groupsystem | −0.026 | 0.17 | 55 | −0.15 | 0.88 | |
| rateLow | −0.18 | 0.05 | 3.8e+05 | −3.6 | 0.00027 | *** |
| rateHigh | −0.097 | 0.051 | 3.9e+05 | −1.9 | 0.057 | . |
| PrepredfactorBetter | −0.25 | 0.056 | 3.8e+05 | −4.4 | 9.2e−06 | *** |
| PrepredfactorWorse | −0.2 | 0.068 | 3.6e+05 | −3 | 0.0028 | ** |
| accuracyerror:sag | 0.38 | 0.15 | 3.9e+05 | 2.5 | 0.011 | * |
| accuracyerror:groupsystem | −0.089 | 0.25 | 67 | −0.36 | 0.72 | |
| sag:groupsystem | 0.06 | 0.093 | 3.9e+05 | 0.65 | 0.52 | |
| accuracyerror:rateLow | 0.26 | 0.12 | 3.8e+05 | 2.1 | 0.035 | * |
| accuracyerror:rateHigh | 0.42 | 0.1 | 3.8e+05 | 4.1 | 4.2e−05 | *** |
| sag:rateLow | 0.16 | 0.095 | 3.9e+05 | 1.7 | 0.096 | . |
| sag:rateHigh | −0.057 | 0.099 | 3.9e+05 | −0.58 | 0.56 | |
| groupsystem:rateLow | 0.11 | 0.069 | 3.8e+05 | 1.7 | 0.098 | . |
| groupsystem:rateHigh | 0.44 | 0.071 | 3.9e+05 | 6.2 | 7.3e−10 | *** |
| accuracyerror:PrepredfactorBetter | 0.74 | 0.12 | 3.4e+05 | 6.1 | 1.1e−09 | *** |
| accuracyerror:PrepredfactorWorse | 0.23 | 0.15 | 3.1e+05 | 1.6 | 0.12 | |
| sag:PrepredfactorBetter | −0.68 | 0.1 | 3.9e+05 | −6.5 | 8.8e−11 | *** |
| sag:PrepredfactorWorse | 0.33 | 0.13 | 3.9e+05 | 2.6 | 0.0093 | ** |
| groupsystem:PrepredfactorBetter | 0.3 | 0.08 | 3.7e+05 | 3.8 | 0.00016 | *** |
| groupsystem:PrepredfactorWorse | 0.61 | 0.099 | 3.6e+05 | 6.1 | 8.4e−10 | *** |
| rateLow:PrepredfactorBetter | 0.22 | 0.074 | 3.8e+05 | 3 | 0.0029 | ** |
| rateHigh:PrepredfactorBetter | 0.4 | 0.086 | 3.8e+05 | 4.6 | 3.6e−06 | *** |
| rateLow:PrepredfactorWorse | 0.27 | 0.11 | 3.7e+05 | 2.5 | 0.014 | * |
| rateHigh:PrepredfactorWorse | 0.21 | 0.091 | 3.7e+05 | 2.3 | 0.022 | * |
| accuracyerror:sag:groupsystem | −1.7 | 0.2 | 3.9e+05 | −8.4 | 3.5e−17 | *** |
| accuracyerror:sag:rateLow | −1 | 0.24 | 3.9e+05 | −4.3 | 1.8e−05 | *** |
| accuracyerror:sag:rateHigh | −0.21 | 0.2 | 3.9e+05 | −1.1 | 0.28 | |
| accuracyerror:groupsystem:rateLow | −0.24 | 0.17 | 3.8e+05 | −1.4 | 0.16 | |
| accuracyerror:groupsystem:rateHigh | −0.31 | 0.14 | 3.8e+05 | −2.1 | 0.032 | * |
| sag:groupsystem:rateLow | −0.012 | 0.13 | 3.9e+05 | −0.091 | 0.93 | |
| sag:groupsystem:rateHigh | −0.38 | 0.14 | 3.9e+05 | −2.7 | 0.0063 | ** |
| accuracyerror:sag:PrepredfactorBetter | 0.3 | 0.23 | 3.9e+05 | 1.3 | 0.2 | |
| accuracyerror:sag:PrepredfactorWorse | −1 | 0.27 | 3.9e+05 | −3.7 | 0.00023 | *** |
| accuracyerror:groupsystem:PrepredfactorBetter | −0.83 | 0.17 | 3.2e+05 | −4.8 | 1.8e−06 | *** |
| accuracyerror:groupsystem:PrepredfactorWorse | −0.46 | 0.22 | 3e+05 | −2.1 | 0.032 | * |
| sag:groupsystem:PrepredfactorBetter | −0.19 | 0.15 | 3.9e+05 | −1.3 | 0.2 | |
| sag:groupsystem:PrepredfactorWorse | −1.6 | 0.18 | 3.9e+05 | −8.8 | 1.4e−18 | *** |
| accuracyerror:rateLow:PrepredfactorBetter | −0.63 | 0.18 | 3.7e+05 | −3.5 | 0.0005 | *** |
| accuracyerror:rateHigh:PrepredfactorBetter | −0.93 | 0.17 | 3.6e+05 | −5.4 | 6.5e−08 | *** |
| accuracyerror:rateLow:PrepredfactorWorse | 0.25 | 0.27 | 3.6e+05 | 0.92 | 0.36 | |
| accuracyerror:rateHigh:PrepredfactorWorse | −0.67 | 0.19 | 3.2e+05 | −3.6 | 0.00033 | *** |
| sag:rateLow:PrepredfactorBetter | 0.17 | 0.14 | 3.9e+05 | 1.2 | 0.23 | |
| sag:rateHigh:PrepredfactorBetter | 1.4 | 0.16 | 3.9e+05 | 8.7 | 3.4e−18 | *** |
| sag:rateLow:PrepredfactorWorse | 0.13 | 0.2 | 3.9e+05 | 0.64 | 0.52 | |
| sag:rateHigh:PrepredfactorWorse | −0.39 | 0.17 | 3.9e+05 | −2.3 | 0.021 | * |
| groupsystem:rateLow:PrepredfactorBetter | −0.2 | 0.1 | 3.8e+05 | −1.9 | 0.058 | . |
| groupsystem:rateHigh:PrepredfactorBetter | −0.65 | 0.12 | 3.8e+05 | −5.4 | 8.4e−08 | *** |
| groupsystem:rateLow:PrepredfactorWorse | −0.56 | 0.15 | 3.6e+05 | −3.6 | 0.0003 | *** |
| groupsystem:rateHigh:PrepredfactorWorse | −0.65 | 0.13 | 3.7e+05 | −5 | 6.2e−07 | *** |
| accuracyerror:sag:groupsystem:rateLow | 1.6 | 0.33 | 3.9e+05 | 4.8 | 2e−06 | *** |
| accuracyerror:sag:groupsystem:rateHigh | 0.89 | 0.28 | 3.9e+05 | 3.2 | 0.0013 | ** |
| accuracyerror:sag:groupsystem:PrepredfactorBetter | 0.39 | 0.32 | 3.9e+05 | 1.2 | 0.23 | |
| accuracyerror:sag:groupsystem:PrepredfactorWorse | 0.44 | 0.4 | 3.9e+05 | 1.1 | 0.27 | |
| accuracyerror:sag:rateLow:PrepredfactorBetter | 0.93 | 0.34 | 3.9e+05 | 2.7 | 0.0068 | ** |
| accuracyerror:sag:rateHigh:PrepredfactorBetter | −0.3 | 0.33 | 3.9e+05 | −0.92 | 0.36 | |
| accuracyerror:sag:rateLow:PrepredfactorWorse | 1.8 | 0.51 | 3.9e+05 | 3.5 | 0.00044 | *** |
| accuracyerror:sag:rateHigh:PrepredfactorWorse | 1.2 | 0.35 | 3.9e+05 | 3.5 | 0.00047 | *** |
| accuracyerror:groupsystem:rateLow:PrepredfactorBetter | 0.99 | 0.25 | 3.8e+05 | 3.9 | 9.3e−05 | *** |
| accuracyerror:groupsystem:rateHigh:PrepredfactorBetter | 1 | 0.24 | 3.6e+05 | 4.3 | 1.5e−05 | *** |
| accuracyerror:groupsystem:rateLow:PrepredfactorWorse | −0.6 | 0.38 | 3.4e+05 | −1.6 | 0.12 | |
| accuracyerror:groupsystem:rateHigh:PrepredfactorWorse | 0.94 | 0.27 | 3.2e+05 | 3.5 | 0.00045 | *** |
| sag:groupsystem:rateLow:PrepredfactorBetter | 0.32 | 0.2 | 3.9e+05 | 1.6 | 0.1 | |
| sag:groupsystem:rateHigh:PrepredfactorBetter | −0.69 | 0.23 | 3.9e+05 | −3 | 0.0028 | ** |
| sag:groupsystem:rateLow:PrepredfactorWorse | 0.014 | 0.28 | 3.9e+05 | 0.048 | 0.96 | |
| sag:groupsystem:rateHigh:PrepredfactorWorse | 1.5 | 0.24 | 3.9e+05 | 6.1 | 1.1e−09 | *** |
| accuracyerror:sag:groupsystem:rateLow:PrepredfactorBetter | −1.9 | 0.49 | 3.9e+05 | −3.9 | 8.6e−05 | *** |
| accuracyerror:sag:groupsystem:rateHigh:PrepredfactorBetter | −0.74 | 0.46 | 3.9e+05 | −1.6 | 0.11 | |
| accuracyerror:sag:groupsystem:rateLow:PrepredfactorWorse | −3.9 | 0.72 | 3.9e+05 | −5.4 | 6.7e−08 | *** |
| accuracyerror:sag:groupsystem:rateHigh:PrepredfactorWorse | −0.87 | 0.5 | 3.9e+05 | −1.8 | 0.079 | . |
